# Clonal expansion from standing genetic variation underpins the evolution of an emerging plant pathogen in Australia

**DOI:** 10.1101/2024.12.26.630446

**Authors:** Adam H. Sparks, Dante L. Adorada, Elena Colombi, Lisa A. Kelly, Anthony Young, Noel L. Knight, Niloofar Vaghefi

## Abstract

Pathogens can evolve rapidly, leading to the emergence of novel strains that can overcome commercially deployed host plant resistance. Understanding the genetic and phenotypic diversity and population dynamics of plant pathogens is crucial to inform breeding programs targeting resistance. Tan spot, otherwise known as bacterial wilt, caused by *Curtobacterium flaccumfaciens* pv. *flaccumfaciens*, is a destructive disease of mungbean in Australia. Since the 1990s, several mungbean cultivars with partial resistance to tan spot have been released; however, cultivars initially rated as moderately resistant were later rated as moderately susceptible to tan spot. This study investigated the genetic and phenotypic diversity and temporal evolutionary dynamics of *C. flaccumfaciens* pv. *flaccumfaciens* in Australian mungbean fields. Whole-genome sequencing of 119 isolates collected from mungbean and other legumes (1986–2019) enabled analyses of pathogen evolution in Australia and in a global context. Results revealed that clonal expansion from standing genetic variation, rather than introduction of novel genotypes, has driven the evolution of this pathogen in Australia since the 1980s. Glasshouse trials confirmed the role of a plasmid in pathogenicity of *C. flaccumfaciens* pv. *flaccumfaciens* but found no significant differences in aggressiveness between clonal lineages. This study underscores the importance of understanding pathogen diversity and aggressiveness to guide breeding programs for effective and durable disease resistance and highlights the need for further studies on the competitive fitness of isolates and characterisation of private alleles linked to dominant clonal lineages.

**Significance:** Tan spot is an increasingly significant pathogen affecting beans and mungbean worldwide, with a broad host range and quarantine importance in many countries underscoring the need for effective breeding strategies to manage the disease. Our research provides insights into genetic and phenotypic diversity of this important plant pathogen and temporal changes in its population structure in Australia, which are crucial for developing informed breeding strategies and managing the spread and impact of the disease. This work also underscores the critical need for robust, validated screening protocols and selection of aggressive, locally representative isolates for resistance breeding to ensure accurate assessment of resistance in crops for effective and durable resistance.

## Introduction

Intensive agricultural ecosystems are characterised by high-density monocultures of genetically uniform crops. These promote evolution of existing pathogens into more aggressive and highly adapted populations and can favour the emergence of novel crop pathogens via hybridisation or host jumps. Monocultures and reduced environmental heterogeneity in agricultural ecosystems result in pathogen populations with large effective sizes, thus, a large number of mutations, which are under constant selection pressure from the environment, host crop, and management practices (Stukenbrock & McDonald 2008; McDonald & Stukenbrock 2016). Other important factors determining adaptive evolution of crop pathogens are the amount of standing genetic diversity in pathogen populations as well as the introduction of new genetic variation via migration or horizontal gene transfer, on which selection can act (Mable 2019).

Accelerated evolution is most apparent in pathogen populations capable of clonal reproduction, where more adapted clonal lineages (generated via mutation or recombination) are asexually propagated and rapidly replace less adapted lineages. This can be augmented via natural and human-mediated movement of inoculum, leading to destructive crop disease epidemics (Goodwin et al. 1994; Li et al. 2012; Vaghefi et al. 2015, 2017). Understanding temporal changes in pathogen populations and the underlying factors promoting their adaptive evolution is critical to effective and sustainable disease management and breeding for resistance. This may be of particular importance for pathogens with the ability to cause disease on multiple crops, which have adapted to an extended geographic and host range, posing great risk to crop production and biosecurity (McDonald & Stukenbrock 2016; Castillo et al. 2021).

The bacterial species *Curtobacterium flaccumfaciens* pv. *flaccumfaciens* is a seedborne, vascular pathogen of legumes. It can infect a wide range of crops, including adzuki bean (*Vigna angularis*), black gram (*V. mungo*), common bean (*Phaseolus vulgaris*), cowpea (*V. unguiculata*), lima bean (*Phaseolus lunatus*), mungbean (*V. radiata*), soybean (*Glycine max*), and sunflower, among others (Osdaghi et al. 2020; Tarakanov et al. 2023; Pilik et al. 2023). It can also survive epiphytically on several non-host crops such as eggplant, maize, pepper, tomato, and wheat (Osdaghi et al. 2018a; Gonçalves et al. 2019).

This pathogen was first discovered in the USA on dry beans (Hedges 1922) and it has since been identified in many legume-producing regions (Osdaghi et al. 2015, 2020; Evseev et al. 2022a). In Europe, the Middle East, and the Americas, *C. flaccumfaciens* pv. *flaccumfaciens* is known as the cause of bacterial wilt of dry bean and soybean (Harveson & Schwartz 2007) while, in Australia, it is mainly known as the cause of tan spot on mungbean, a summer legume grown predominantly in northern New South Wales and Queensland.

In Australia, the release of high-yielding and non-shattering mungbean cultivars Berken and Celera promoted extensive production of mungbean as a grain crop in the 1970s (Kelly et al. 2021). Tan spot was first detected in central Queensland in mungbean fields in 1984, and later in southern Queensland on mungbean and in northern New South Wales on cowpea, in 1986 (Wood & Easdown 1990). Subsequently, tan spot became one of the most destructive diseases of mungbean in Australia, with incidences of up to 90% recorded in some fields (Wood & Easdown 1990; Vaghefi et al. 2019).

Previous *C. flaccumfaciens* pv. *flaccumfaciens* population genetic analyses using amplified fragment length polymorphisms, repetitive element sequence-based PCR, pulse field gel electrophoresis markers, and multi-locus phylogenetic analyses revealed that populations in different parts of the world are genetically heterogeneous, with no clear association of genetically distinct lineages with host of origin, geographical location, or pathogenicity (Agarkova et al. 2012; Gonçalves et al. 2019). A high level of phenotypic diversity was also detected in populations worldwide, with different colony types producing yellow-, orange-, red-, pink- and purple-pigmented colonies (Harveson & Schwartz 2007; Osdaghi & Lak 2015; Osdaghi et al. 2016). In Australia, however, only the yellow-pigmented type has been detected thus far. This, together with the hypothesis that the American High Plains is the centre of origin of this pathogen (Agarkova et al. 2012), suggests that *C. flaccumfaciens* pv. *flaccumfaciens* may be an introduced, invasive species in Australia.

Breeding for tan spot resistance has been central to the national mungbean improvement program in Australia, which has generated multiple mungbean cultivars with partial resistance to tan spot since the 1990s. However, mungbean cultivars rated as moderately resistant at the time of their commercial release, *e.g.*, cultivars Celera and Crystal (Gentry & Cumming 2008; Gentry 2010), were subsequently rated as moderately susceptible to tan spot after a few years (Douglas & Cumming 2013, 2014; GRDC 2017). The factors underlying these shifts in the resistance ratings of Australian mungbean cultivars remain unclear.

This study is the first population genetic study of *C. flaccumfaciens* pv. *flaccumfaciens* in Australia. Our objectives were to investigate the origin of contemporary *C. flaccumfaciens* pv. *flaccumfaciens* genotypes responsible for tan spot epidemics in Australian mungbean fields; examine the evolutionary relationships among Australian and overseas isolates to detect potential introductions; and explore the temporal evolution and potential adaptation of the pathogen since its emergence in Australian mungbean fields in the 1980s. To achieve this, we conducted whole genome sequencing and population genomic analyses of 119 historical and contemporary *C. flaccumfaciens* pv. *flaccumfaciens* isolates collected from mungbean and other legumes from 1986 to 2019 in Australia. We examined temporal changes and signatures of selection and assessed the evolutionary relationships of Australian *C. flaccumfaciens* pv. *flaccumfaciens* isolates with those from the USA and Eurasia to identify potential introductions. Additionally, a subset of isolates from different clonal lineages and with varying plasmid content was evaluated for aggressiveness on mungbean and other crops.

## Results

### Genotypic diversity and spatial and temporal distribution of multi-locus genotypes (MLGs)

A total of 130 genomic DNA samples (119 unique isolates and 11 within-plate and between-plate replicate DNA samples) were genotyped using Illumina paired-end sequencing (average 45× coverage) and reference-based variant-calling. Removal of unreliable sites that carried different alleles between the replicated DNA samples and filtering the single nucleotide variations for a minimum allele frequency of 0.05 and no missing data, resulted in identification of 43,974 polymorphic sites (Single Nucleotide Polymorphisms; SNPs) on the main chromosome (core SNPs) and 99 sites on the plasmid (plasmid SNPs) in 117 *C. flaccumfaciens* pv. *flaccumfaciens* isolates (Supplementary Table S1). Two isolates were removed from the data set due to greater than 20% missing data.

The collection of 117 isolates included 97 contemporary isolates collected from 2014 to 2019 across 26 fields in New South Wales (NSW) and Queensland (Qld), and 21 historical isolates collected in 1986 and 1987, when *C. flaccumfaciens* pv. *flaccumfaciens* was first reported in mungbean fields in Australia (obtained from NSW Plant Pathology and Mycology Herbarium, New South Wales Department of Primary Industries [DAR Culture Collection, Orange, New South Wales]) (Figure 1; Supplementary Table S1). All isolates were confirmed as *C. flaccumfaciens* pv*. flaccumfaciens* through PCR and Sanger sequencing of the 16S region, which was identical to that of the ex-type isolate, LMG 3645 (NCBI accession no. AJ312209). All isolates belonged to the yellow-pigmented colony type.

**Fig. 1.**
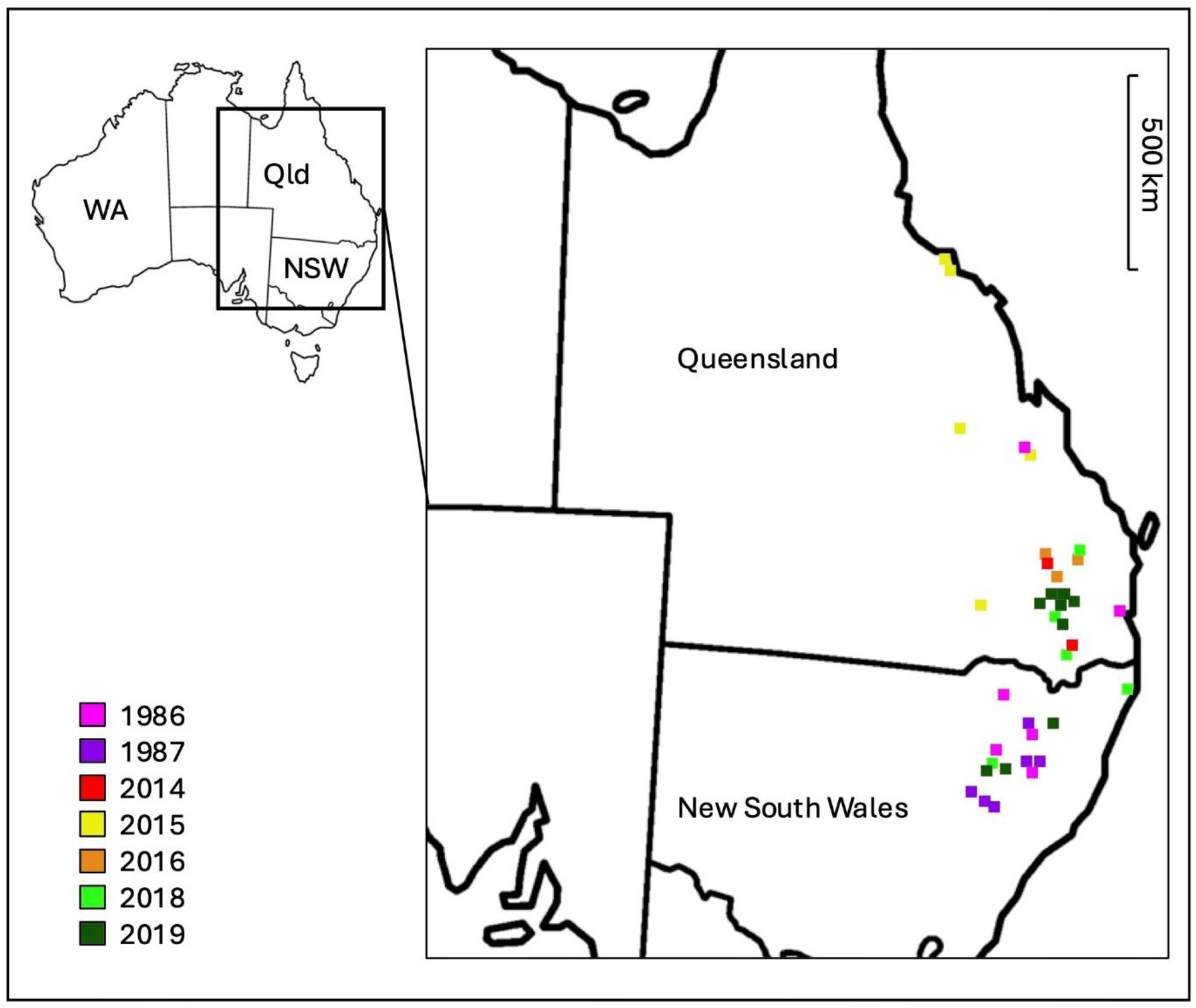
Geographical origins of 119 *Curtobacterium flaccumfaciens* pv. *flaccumfaciens* isolates from mungbean growing regions in New South Wales (NSW) and Queensland (Qld), Australia, sequenced in this study. Colours correspond to year of collection.

The 43,974 core SNPs were used for delineation of unique multi-locus genotypes (MLGs) and multi-locus linages (MLLs) and subsequent genetic diversity analyses. In clonal populations, genotypic diversity may be overestimated using whole genome SNP data due to the accumulation of mutations, which will result in unique MLGs that are differentiated based on only a handful of loci within clonal lineages (Kamvar et al. 2015; Milgroom 2017; Vaghefi et al. 2017). Following inspection of the dataset for pairwise genetic distance, a total of 66 MLGs were detected, 20 of which differed at only one to seven SNP loci out of 43,974 sites (Supplementary Figure S1). Therefore, clonal boundaries were delineated based on a defined genetic distance threshold estimated using *cutoff_predictor* in *poppr* (Kamvar et al. 2014, 2015) and MLGs that differed in only one to seven SNPs were collapsed into the same MLL, resulting in identification of 40 MLLs in the total of 117 isolates (Supplementary Table S1). This indicated a clonal fraction of 0.66 (N-number of MLLs)/N) where N is the total number of individuals, and unbiased Simpson’s (Simpson 1949) complement index of genotypic diversity of 0.89 for the Australian population (*n* = 117) (Supplementary Table S2). The term population here is used to refer to a set of isolates defined under certain geographical or temporal criteria, *e.g.*, collected from a location (State, field) or in the same year.

Populations from the two intensively sampled fields had similar genotypic diversity, as indicated by expected number of MLLs estimated after rarefaction (Supplementary Table S3). While in some cases only one MLL was isolated per leaf, up to three different MLLs were retrieved from different lesions on the same leaf (Supplementary Table S1). Plotting total genotypic diversity by sample size showed a plateau was reached for Field 42, indicating that sampling 18 isolates per field was enough to capture > 80% of the total genotypic diversity (Supplementary Figure S2).

Analyses of temporal and geographical origins of isolates showed that several MLGs and MLLs were shared among different years and locations (Supplementary Figures S3 and S4). MLL35 (including MLGs 35, 36, 37, 38, 39, 40, and 41) was the most frequent (*n* = 34), which was re-isolated across four years (2015, 2016, 2018, and 2019) and from 11 locations in NSW and Qld. Private allele analyses detected 10 SNPs (two low impact, seven moderate impacts, and one high impact) to be associated with MLL35. Seven and one of these SNPs were predicted to result in missense and nonsense mutations, respectively, affecting predicted transporter proteins and transcriptional regulators (Supplementary Table S4).

The second most frequent MLL was MLL43 (*n* = 11) (MLGs 42, 43, and 44), occurring in 1986 and 2018, and in five locations in NSW and Qld. MLL21 and MLL27 recurred across hosts, as both were isolated from black gram and cowpea. Even when MLGs were analysed (splitting MLLs into different genotypes differing by 1 to 4 SNPs), similar results were obtained with multiple MLGs recurring across years and locations (Supplementary Figure S4). No private alleles were detected for MLL21, MLL27 and MLL43. The Mantel test for the correlation between genetic distance decay and geographic distance did not detect a significant relationship between MLGs and geographical location (*r* = 0.0003454, *p* value = 0.542). No statistically significant correlation was detected between geographic and genetic distance considering the effect of time in a partial Mantel test (*r* = -0.04338, *p* value = 0.722).

### Population structure and temporal differentiation of *Curtobacterium flaccumfaciens* pv. *flaccumfaciens* in Australia

Analysis of population structure without *a priori* assumption of populations using STRUCTURE (Pritchard et al. 2000) detected six clusters/subpopulations within the *Curtobacterium flaccumfaciens* pv. *flaccumfaciens* population in Australia (Figures 2a and 2b). Ten admixed isolates could not be confidently assigned to any one cluster based on likelihood probability of > 0.6 (Supplementary Table S1). Statistically significant differences (Fisher’s exact test *p* < 0.05) were detected among the frequencies of isolates in clusters between the historical (1986 and 1987) and contemporary (2014 to 2019) populations. Post-hoc examination of adjusted standardized residuals indicated that the frequency of isolates belonging to Clusters 1, 3, 4, and 6 varied significantly between the two populations. Isolates belonging to Clusters 1, 2, and 5 occurred in both the historical and contemporary populations (Figure 2c); the frequency of isolates belonging to Cluster 1 was significantly greater in the contemporary population while the frequency of isolates belonging to Clusters 2 and 5 was not significantly different. Cluster 3, the most frequently sampled cluster in the historical population, was not detected in the contemporary population, while Clusters 4 and 6 were only identified in the contemporary population (Figure 2c). Cluster 1 included 68 isolates obtained exclusively from mungbean across 22 locations from 1986 to 2019 and was the most frequently sampled cluster in the contemporary population (obtained from 36% of sampled locations in 1986/87, while occurring in > 70% of the locations sampled from 2014 to 2019). Clusters 4 and 6 were also only associated with mungbean, whereas Clusters 2 and 5 were isolated from multiple hosts including mungbean. Cluster 3 was only associated with hosts other than mungbean.

**Fig. 2.**
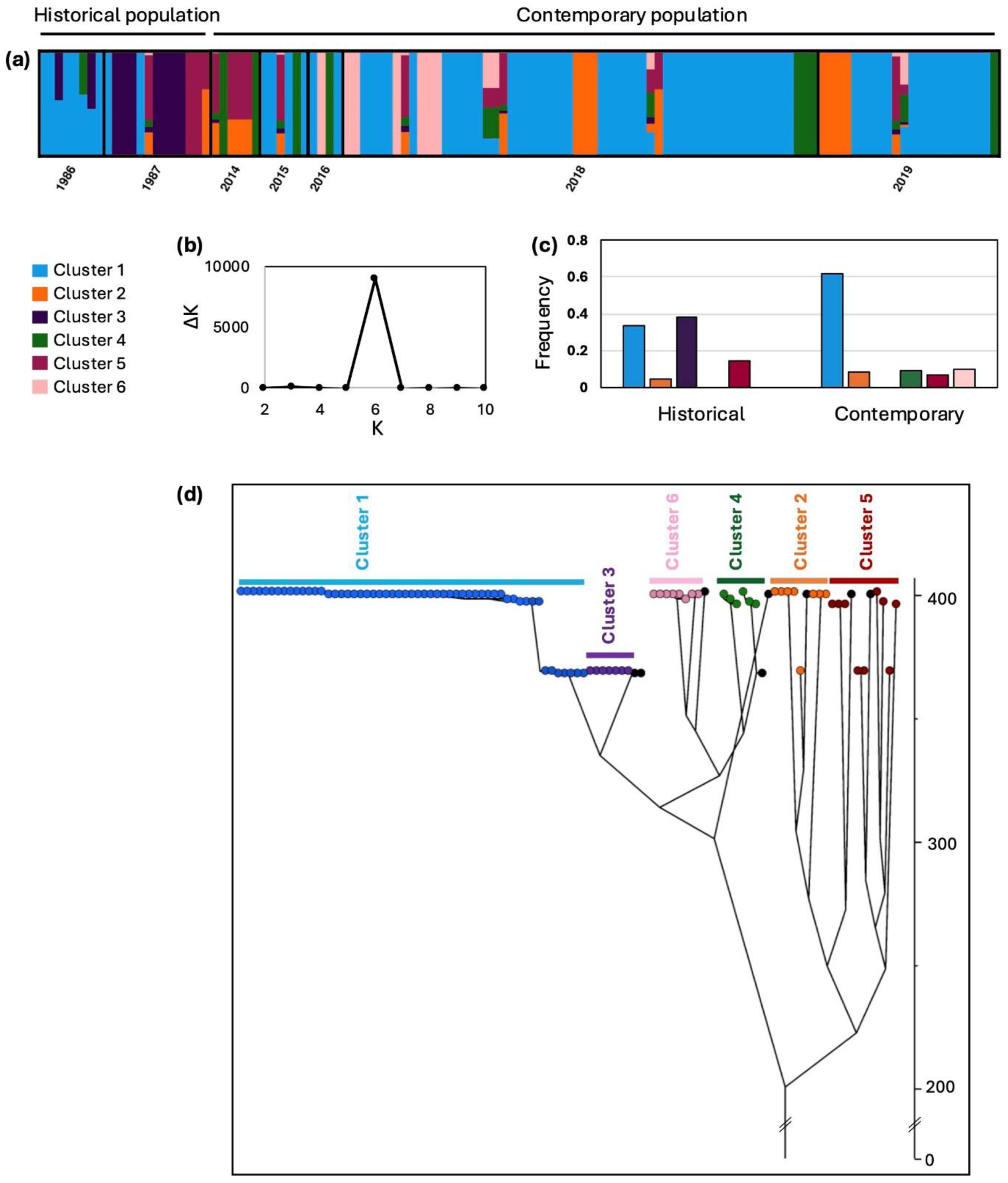
**(a)** Assignment of *Curtobacterium flaccumfaciens* pv. *flaccumfaciens* isolates collected from 1986 to 2019 from various host crops in Australia to six clusters detected through Bayesian clustering analysis of 30,035 single nucleotide polymorphisms with predicted neutral or low impact. Each bar represents one isolate and the bar height indicates estimated membership fraction of each individual in the inferred clusters **(b)** ΔK calculated as ΔK = m|L′′(K)|/s[L(K)] based on Evanno et al. (2005), showing the peak at K = 6; K represents the inferred number of clusters and ΔK peaks at the most likely value of K **(c)** Frequency distribution of the isolates belonging to the six clusters identified via STRUCTURE version 2.3.4 (Pritchard et al. 2000), within the historical (1986 and 87) and contemporary (2014 to 2019) populations **(d)** Dated phylogeny of *C. flaccumfaciens* pv. *flaccumfaciens* isolates inferred using Bayesian phylodynamic analysis in BEAST (Bouckaert et al. 2019) after clone-correction. Numbers across the Y axis denote years since divergence. Branch tips are colour-coded by the clusters detected by STRUCTURE analysis. Admixed isolates, which could not be assigned to any clusters with > 0.6 probability, are shown in black.

A similar division of the Australian *C. flaccumfaciens* pv. *flaccumfaciens* population into six subpopulations was observed using Discriminant Analysis of Principal Components (DAPC) in *adegenet* (Jombart 2008) using the same set of SNPs (Supplementary Fig. S5). Individuals in each of the six subpopulations in the DAPC analysis corresponded to the six clusters identified using STRUCTURE. Repeating the STRUCTURE and DAPC analyses using all SNPs (including the SNPs with predicted high and moderate impact) resulted in detection of identical number of clusters and assignment of isolates (*data not shown*).

### Phylodynamic analysis

Bayesian phylodynamic analysis using BEAST after clone-correction (Figure 2d) illustrated the clonal expansion of Cluster 1. Here we use the term “expansion” as an increase in the number of clones within a clonal lineage, which could be either due a selective advantage or simple founder effect (Helekal et al. 2022). The very short branch lengths within Cluster 1 indicates small genetic distances among isolates. Although the estimated phylogeny is dated based on year of collection, it should be noted that the Bayesian analysis is not used to infer exact divergence times but to infer relative evolutionary histories as sampling dates do not necessarily correspond to introduction dates. Analyses conducted with different priors yielded consistent results. The result based on a strict molecular clock and coalescent exponential growth is shown in Figure 2d.

### Outlier analyses for detection of loci potentially under selection

Outlier analysis using fixation index (*F*_st_) comparisons of SNPs between the historical and contemporary populations identified 30 SNPs potentially under selection above the 99% confidence level (1% FDR; false discovery rate defined as the proportion of incorrectly identified outliers (false positives) among all the detected outliers). These SNPs were associated with 18 synonymous variations in several genes with predicted roles in metabolic and signalling pathways as well as five intergenic variations and seven missense variations. Three of the missense variations were associated with genes with predicted functions in cell division while four were located on loci encoding hypothetical proteins of unknown function (Supplementary Table S5). Outlier analysis using BayeScan (Foll & Gaggiotti 2008), which is also based on allele frequencies but takes population structure into account, did not detect any outlier loci based on pairwise comparison of the historical and contemporary populations even at false discovery rates as high as 5% or 10%.

### Australian *Curtobacterium flaccumfaciens* pv. *flaccumfaciens* population within a global context

Genome assemblies of 27 non-Australian *C. flaccumfaciens* pv. *flaccumfaciens* isolates available in December 2023 were retrieved from the national centre for biotechnology information (NCBI) database to be used as reference genomes (Supplementary Table S6). An initial phylogeny using 1,669 single-copy core genes identified two clades of non-Australian isolates which were separated from the rest of the collection. The two clades each consisted of four isolates, CFBP 3417, CFBP 3422, Carlos3, and Carlos7 in one clade; and CFBP 8818, CFBP 8819, CFBP 8823, and CFBP 8824 in another (Supplementary Figure S6). These had an average pairwise nucleotide identity (ANI) < 95% compared to the *C. flaccumfaciens* pv. *flaccumfaciens* type strain LMG 3645 (= CFBP3418 = NCPPB 1446) (Collins & Jones 1983). Since 95% is defined as the ANI threshold used to demarcate species boundaries in prokaryotes (Jain et al. 2018), the aforementioned isolates were excluded from further analyses, except for Carlos 7, which was used as an outgroup.

In order to place the Australian *C. flaccumfaciens* pv. *flaccumfaciens* isolates in a global context, whole genome sequence data of the remaining 19 isolates from Eurasia and the USA and a subset of Australian isolates (at least one representative per clone was included) were used to produce a phylogeny using 1,835 core genes (a 1,611,342 bp gap-free alignment) (Figure 3 and Supplementary Figure S7). The phylogenetic network (Figure 3) and maximum likelihood tree (Supplementary Figure S7) were in general agreement, with the Australian MLGs representing STRUCTURE Clusters 1, 3, 4 and 6 grouping separately from the overseas isolates while isolates from STRUCTURE Clusters 2 and 5 were interspersed among isolates from different hosts and countries. Based on the sequences used in this study, it appears that the genotypes represented in STRUCTURE Clusters 1, 3, 4 and 6 are specific to Australia, but the lack of reference isolates from overseas constitutes a possible sampling-based bias.

**Fig. 3.**
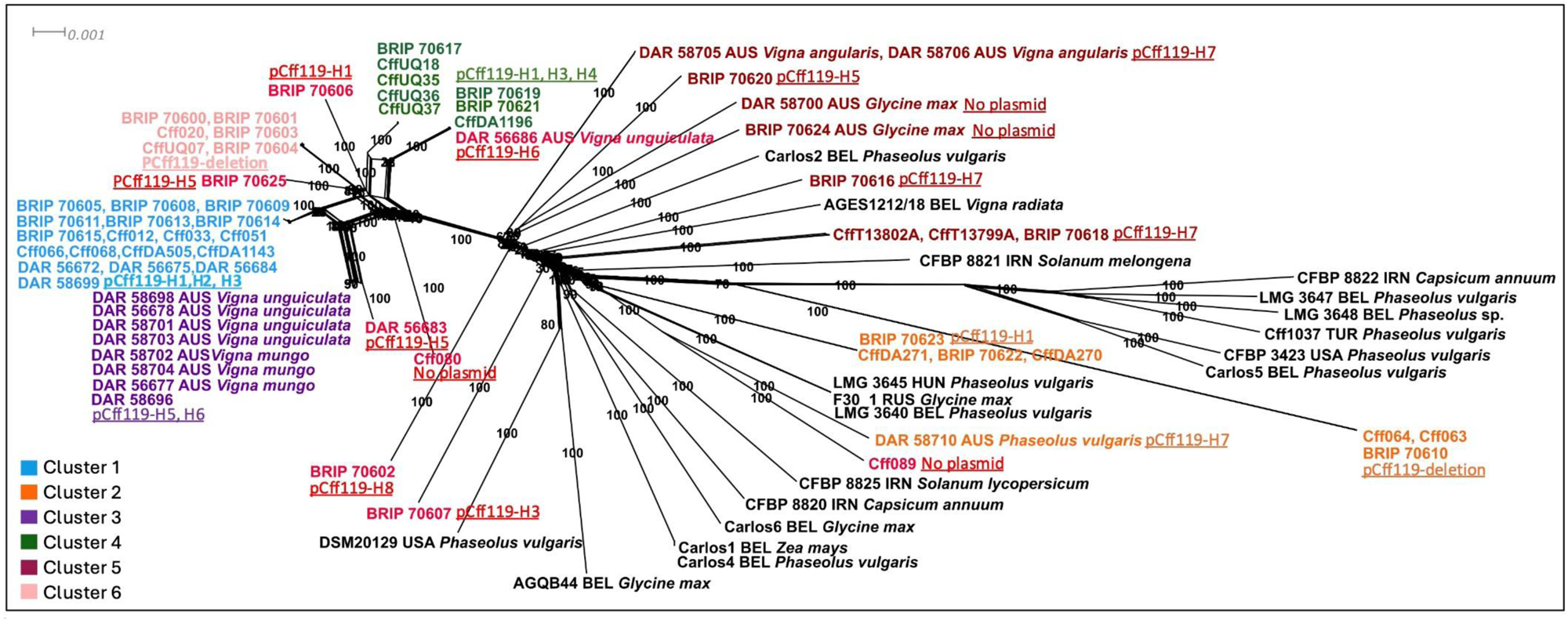
Phylogenetic relationships among a global population of *Curtobacterium flaccumfaciens* pv*. flaccumfaciens* based on nucleotide sequences of 1,835 core genes constructed in Splitstree (Huson & Bryant 2006). Tip colours correspond to the clusters detected by the software STRUCTURE (Pritchard et al. 2000) (Fig. 2). Admixed isolates which could not be assigned confidently to a cluster with probability of > 0.60 are indicated in red. Plasmid haplotypes for Australian isolates occurring in each cluster are underlined and indicated in the same colour (plasmid haplotypes defined in Fig. 4). Bootstrap support values above 80% are shown next to the branches. The scale bar represents the expected nucleotide changes per site.

### Movement of a plasmid within the Australian *Curtobacterium flaccumfaciens* pv. *flaccumfaciens* population and its role in pathogenicity

The singular linear plasmid named pCff119 (previously reported by Vaghefi et al. 2021) was identified in the genome of 113 of the historical and contemporary *C. flaccumfaciens* pv. *flaccumfaciens* isolates in Australia. Four isolates, including historical isolate DAR 58700, and the contemporary isolates Cff080, Cff089, and BRIP 70624, did not contain the pCff119 or any other plasmids. Of the 113 isolates carrying the pCff119 plasmid, nine (belonging to five MLLs) carried a truncated version of pCff119 (denoted as the pCff-deletion haplotype). This truncated version of pCff119 lacks an 8 kb region from the middle of pCff119 (CP074440.1 coordinates: 25,286 to 33,515) and has previously been reported in Australia and Turkey (O’Leary & Gilbertson 2020; Vaghefi et al. 2021). These included all isolates from Cluster 6 and three isolates from Cluster 2.

The remaining 104 Australian isolates had a complete version of pCff119, with eight unique plasmid haplotypes identified based on 99 plasmid SNPs (Figure 4). The plasmid haplotype pCff119-H1 had the highest frequency in the contemporary population and also occurred in 1986. Haplotypes pCff119-H7 and pCff119-H8 had an accumulation of 76 and 75 SNPs, respectively (Figure 4). The complete pCff1 plasmid, which was reported to have an additional 27 kb region making it a circular plasmid, and pCff2 and pCff3 constructs previously reported in *C. flaccumfaciens* pv. *flaccumfaciens* (Chen et al. 2021) were not detected in any of the Australian isolates.

**Fig. 4.**
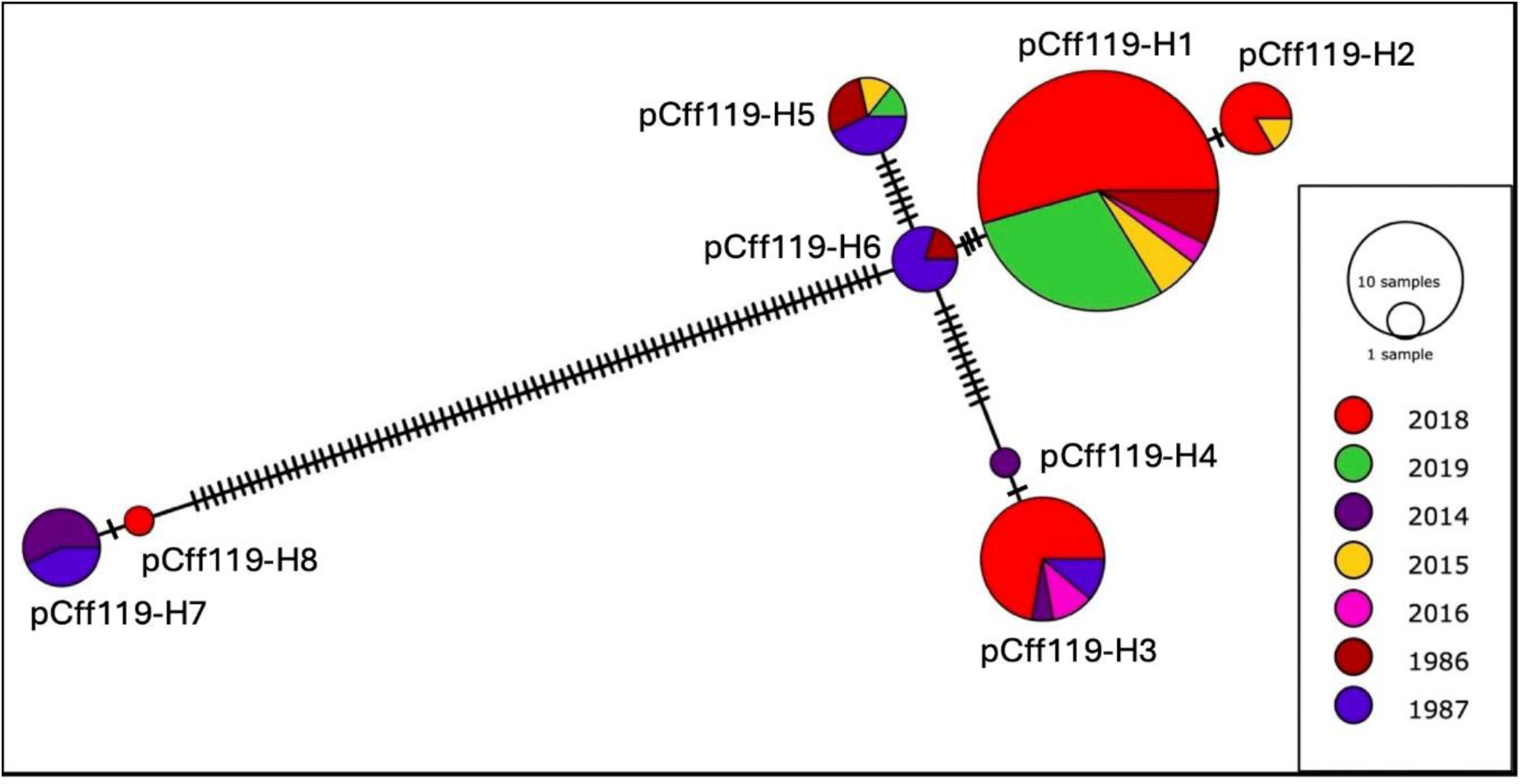
Minimum spanning haplotype network of plasmid haplotypes of 104 Australian *Curtobacterium flaccumfaciens* pv. *flaccumfaciens* isolates with a complete pCff119 plasmid (pCff-deletion haplotype is not included). The network was generated based on 99 single nucleotide polymorphisms located on pCff119 plasmid of *C. flaccumfaciens* pv. *flaccumfaciens* isolates collected from 1986 to 2019 in Australia. Australian plasmid haplotypes (pCff119-H) were detected by the software POPART version 1.7 (Leigh & Bryant 2015). The size of the circles corresponds the total number of samples belonging to each haplotype. Hatch marks on branches indicate the number of mutations. Colours correspond to year of collection.

Incongruence of the plasmid haplotypes and genetic clusters (detected using the SNPs on the core genome) was observed, with the same plasmid haplotypes occurring in several distinct clades in the *C. flaccumfaciens* pv. *flaccumfaciens* phylogeny (Figure 3). For example, plasmid haplotype pCff119-H1 occurred in four distinct clades in the Australian population and plasmid pCff-deletion haplotype occurred in Clusters 2 and 6. This suggests horizontal transfer of plasmid haplotypes among clusters.

A similar pattern was also detected in the global population with multiple overseas isolates carrying similar haplotypes to the Australian isolates (Supplementary Figure 7), in line with the horizontal transfer of the plasmid among isolates. For example, isolate AGES1212/18 from *V. radiata* in Belgium also carried a pCff119 plasmid haplotype with an accumulated number of SNPs, similar to three historical and five contemporary isolates from Australia that carried pCff119-H7 and pCff119-H8. The plasmid of these isolates included > 70 SNPs in an approximately 4.7 kb region (CP074440.1 coordinates: 55,130 to 59,849), while fewer than 30 SNPs were distributed across the rest of the plasmid.

Conjugation systems have been shown to have a role in horizontal transfer of mobile elements in bacteria (Francia et al. 2004). None were detected in the *C. flaccumfaciens* pv. *flaccumfaciens* genomes. However, the hidden-Markov models used by CONJscan (Cury et al. 2020) detected the presence of elements associated with a type 4 secretion system (T4SS), including ATPase VirB4 and Ti-plasmid ATPase VirB11, and the T4SS coupling protein TcpA (Abby et al. 2016) in pCff119. Conjugative DNA relaxases, also referred to as ‘MOB’ proteins, are grouped into six MOB families: MOB_F_, MOB_H_, MOB_Q_, MOB_C_, MOB_P_ and MOB_V_. The MOB proteins recognise and nick the origin-of-transfer sequences on the DNA of conjugative elements. This nicking is required for initiating the transfer of single stranded DNA through the conjugative T4SS to the recipient cell (Francia et al. 2004; Abby et al. 2016). The presence of relaxases was evaluated by searching for MOB-family proteins using hidden-Markov models, with a MOB_F_ relaxase found on pCff119 (CP074440.1, locus tag: JG551_003678).

Pathogenicity tests using three isolates which lacked the plasmid (DAR 58700, Cff089, and BRIP 70624) on *Vigna mungo* (cv. Onyx-AU), *V. radiata* (cv. Jade-AU) and *V. unguiculata* (cv. Red Caloona) demonstrated that these isolates were not able to cause disease symptoms on any of the plants. This suggests a key role of this plasmid in pathogenicity of *C. flaccumfaciens* pv. *flaccumfaciens* (Supplementary Figure S8 and Table S7).

### Aggressiveness of *C. flaccumfaciens* pv. *flaccumfaciens* isolates on mungbean

The aggressiveness of 13 *C. flaccumfaciens* pv. *flaccumfaciens* isolates carrying different plasmid haplotypes from different Clusters (as identified by the software STRUCTURE) on mungbean cv. Jade-AU was assessed. The results from the two independent pathogenicity assays were similar in terms of ranking of isolates based on aggressiveness; therefore, results from the two runs were combined (Table 1). The results supported the absence of pathogenicity of isolates BRIP 70624 (Cluster 5) and Cff089 (Cluster 2), which lacked a plasmid. Limited differences in disease severity was detected among the plasmid-carrying isolates, with two isolates from Cluster 2 (BRIP 70610 and BRIP 70623) causing statistically greater disease severity (*p* < 0.05) compared to isolates Cff14074, BRIP 70620 and CffUQ07 from Clusters 1, 5 and 6, respectively (Figure 5; Table 1).

**Fig. 5.**
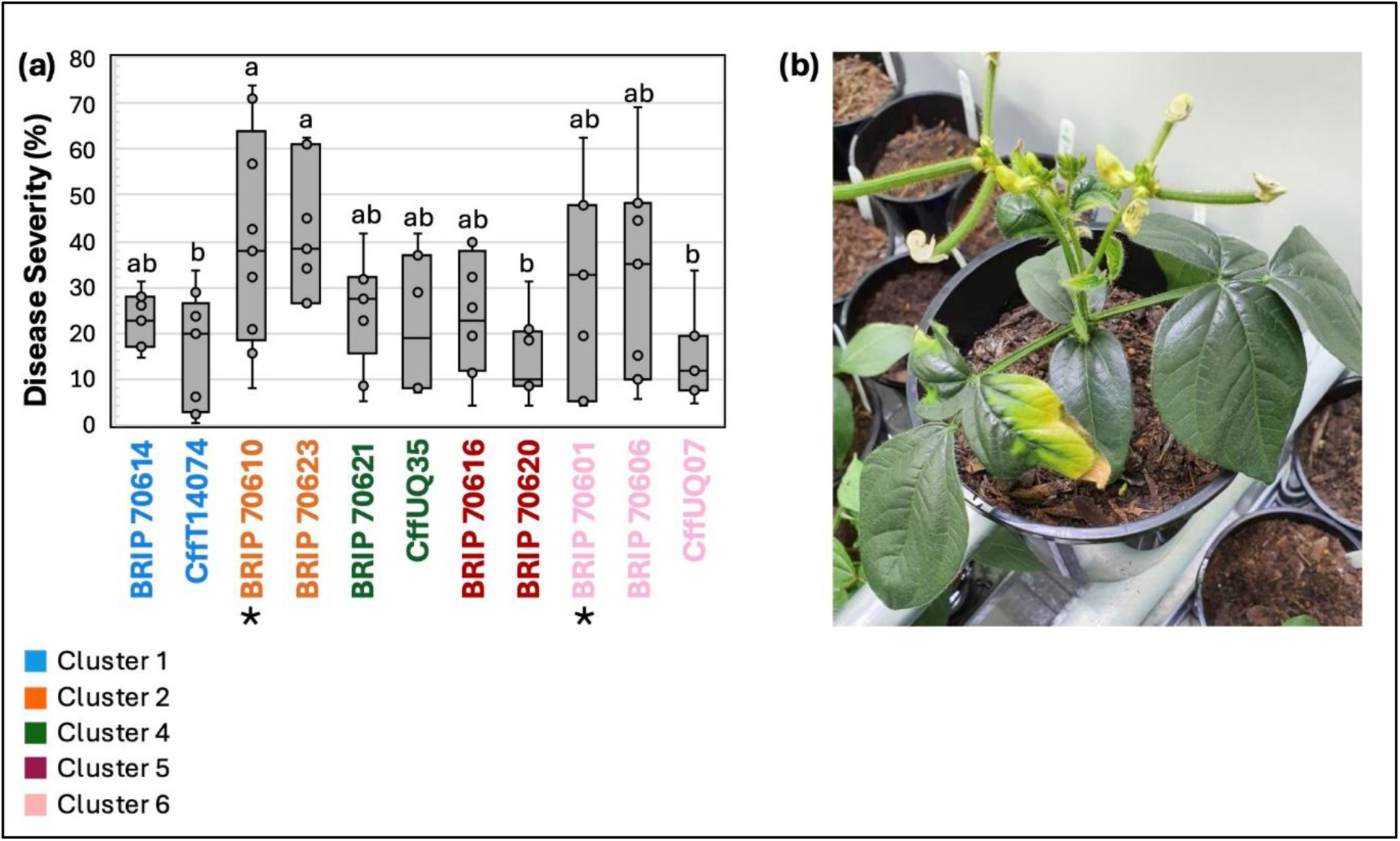
**(a)** Disease severity (percent diseased area) caused by 11 *Curtobacterium flaccumfaciens* pv. *flaccumfaciens* isolates on mungbean cv. Jade-AU. Isolates indicated by an asterisk (*) carry a partial plasmid haplotype (pCff119-deletion). Isolate colours correspond to genetic Clusters identified by the software STRUCTURE (Fig. 2). Different letters on boxplots indicate groups that are significantly different based on pairwise Mann-Whitney U tests, with significance adjusted using Bonferroni correction for multiple comparisons. It should be noted that Kruskal-Wallis test evaluates differences in the median ranks of groups, thus, significant results were detected based on rankings, even if there is overlap in the spread of the data. (b) Tan spot symptoms (leaf chlorosis and necrotic dieback) caused by *C. flaccumfaciens* pv. *flaccumfaciens* on mungbean cv. Jade-AU at 25 days post inoculation.

**Table 1.** Disease severity caused by genetically diverse *Curtobacterium flaccumfaciens* pv. *flaccumfaciens* isolates on *Vigna radiata* cv. Jade-AU in controlled environment trials.

| Cluster<sup>a</sup> | Isolate | Year | Plasmid haplotype<sup>b</sup> | Mean disease severity<sup>c</sup> (%) (*n* = 9) |
| --- | --- | --- | --- | --- |
| Cluster1 | BRIP 70614 | 2018 | pCff119-H1 | 22.4 ab |
| Cluster1 | CffT14074 | 2015 | pCff119-H1 | 15.8 b |
| Cluster2 | BRIP 70623 | 2018 | pCff119-H1 | 39.9 a |
| Cluster2 | BRIP 70610 | 2019 | pCff119-deletion | 41.2 a |
| Cluster2 | Cff089 | 2018 | No plasmid | - |
| Cluster4 | BRIP 70621 | 2015 | pCff119-H1 | 25.4 ab |
| Cluster4 | CffUQ35 | 2018 | pCff119-H3 | 22.1 ab |
| Cluster5 | BRIP 70624 | 2018 | No plasmid | - |
| Cluster5 | BRIP 70616 | 2014 | pCff119-H7 | 21.4 ab |
| Cluster5 | BRIP 70620 | 2015 | pCff119-H5 | 13.9 b |
| Cluster6 | BRIP 70601 | 2018 | pCff119-deletion | 29.2 ab |
| Cluster6 | CffUQ07 | 2018 | pCff119-H3 | 14.1 b |
| Admixed | BRIP 70606 | 2018 | pCff119-H1 | 32.5 ab |

<sup>a</sup> Genetic cluster assigned based on the STRUCTURE analysis depicted in Fig. 2. Isolates from Cluster 3 could not be included due to restrictions imposed by a material transfer agreement not allowing the use of historical isolates for *in vivo* studies.
<sup>b</sup> Haplotype designation was based on 99 single nucleotide polymorphisms in the pCff119 plasmid, as depicted in Fig. 3.
<sup>c</sup> Disease severity was measured as percentage of necrotic and chlorotic leaf area averaged over the first and second trifoliates. Different letter combinations depict statistically significant differences based on Mann-Whitney U tests, with significance adjusted using Bonferroni correction for multiple comparisons ( $p < 0.05$ ).

## Discussion

The *C. flaccumfaciens* pv*. flaccumfaciens* population causing current tan spot outbreaks in Australian mungbean fields originated from outbreaks in the 1980s. A single clonal lineage from the 1980s (Cluster 1) has persisted and expanded clonally, resulting in a large population of closely related MLGs dominating Australian mungbean fields over the past decade. In contrast, another clonal lineage (Cluster 3), which was the most frequent lineage in the 1980s, was not detected in the contemporary population. Notably, all Cluster 3 isolates originated from hosts other than mungbean, and therefore, could still exist in Australian soybean or cowpea fields.

There is limited evidence for the introduction of novel genotypes into Australia, with any such introductions having negligible impact on the contemporary *C. flaccumfaciens* pv*. flaccumfaciens* population in Australian mungbean fields. While several Australian genotypes cluster closely with isolates from Eurasia and the USA, their low frequency suggests that these genotypes are not widely established in Australian mungbean fields. Therefore, although multiple *C. flaccumfaciens* pv*. flaccumfaciens* isolates have potentially been introduced to Australia (either historically or recently), these may have failed to establish in the region due to factors such as poor adaptation to Australian environmental conditions.

Analyses to detect signatures of selection were inconclusive. Pairwise F_st_ analysis identified 30 SNPs potentially under selection between the 1980s and 2019, but BayeScan detected no outlier loci. The differing results may stem from the fundamental differences in the methodologies, assumptions, and sensitivities of the two approaches. F_st_ outlier analysis involves calculating fixation indices for each SNP and identifies loci with unusually high or low values compared to a neutral expectation. Thus, this method assumes that the majority of the genome is evolving neutrally and that outlier loci represent deviations due to selection, without accounting for population structure. BayeScan, on the other hand, uses a Bayesian framework to partition F_st_ into population-specific effects and locus-specific effects, modelling selection explicitly and accounting for population structure and demographic history (Foll & Gaggiotti 2008). The underlying population structure in the Australian *C. flaccumfaciens* pv*. flaccumfaciens* population may have led to false positives in the F_st_ analyses. Dividing populations into clusters for F_st_ analysis was not feasible due to the small number of individuals in each cluster in the historic population.

Evidence continues to highlight the crucial role of plasmid pCff119 in the pathogenicity of *C. flaccumfaciens* pv*. flaccumfaciens*. Vaghefi et al. (2021) identified multiple putative pathogenicity related genes in pCff119, and isolates lacking the plasmid were not pathogenic on mungbean, cowpea or black gram (isolates BRIP 70624, Cff089, DAR 58700) in the current study. These findings align with previous reports indicating that isolates lacking this plasmid failed to cause disease on bean (Osdaghi et al. 2018b, 2022). This suggests that the PCR assay developed for detecting *C. flaccumfaciens* pv*. flaccumfaciens*, which targets a gene encoding a serine protease located on the plasmid (Tegli et al. 2002), may specifically detect pathogenic isolates.

The movement of plasmid pCff119 may confer competitive advantages, potentially through horizontal transfer via a plasmid encoded conjugation system. This is based on the presence of a MOBF relaxase, a T4SS coupling protein, VirB4, and the distribution of plasmids with signature truncation and SNP patterns across the phylogeny of *C. flaccumfaciens* pv*. flaccumfaciens*. It may be hypothesised that: i) pCff119 is responsible for the evolution of pathogenicity in *C. flaccumfaciens* pv*. flaccumfaciens*; ii) after the initial introduction of *C. flaccumfaciens* pv*. flaccumfaciens* in Australia, pCff119 transferred to a local population and mediated the transfer of genes from other sources. For instance, Cff080, the most recent common ancestor of the majority of Australian isolates, lacks the linear plasmid (unfortunately, this isolate has been lost and could not be tested *in planta*). The monophyletic clade including isolates from Clusters 1, 3, 4, and 6, could represent an Australian *C. flaccumfaciens* pv*. flaccumfaciens* population that, after acquiring the plasmid (and associated pathogenicity), increased in frequency in Australian mungbean fields. Further comparative genomic studies of saprophytic and epiphytic *Curtobacterium flaccumfaciens* populations may help test this hypothesis. An example of such phenomenon is seen in the Gram positive *Rhodococcus*, where the evolution of pathogenicity was found to be mediated by the acquisition of a linear plasmid, which transformed *Rhodococcus* from a root hair growth promoting bacterium to a pathogen producing leafy galls in *Nicotiana benthamiana* (Savory et al. 2017). Moreover, the acquisition by locally-adapted bacteria of mobile genetic elements transferred from foreign inocula, which enable the transition from soil dwelling bacteria to plant symbionts, is common in the Australian indigenous population of *Mesorhizobium* (Colombi et al. 2023).

Five out of six Australian isolates sequenced by Vaghefi et al. (2021) carried a single linear plasmid named pCff119 (Vaghefi et al. 2021). In contrast, the PacBio assembly of ICMP 22053, a *C. flaccumfaciens* pv. *flaccumfaciens* isolate from Iran, was reported to carry three circular plasmids pCff1, pCff2 and pCff3 (Chen et al. 2021). The first 119 kb of pCff1 shares 99.9% similarity with pCff119, but pCff1 also contains an additional 27 kb at the 3′ end, resulting in its self-complementarity and circularity (Vaghefi et al. 2021). The additional 27 kb in pCff1, as well as the two additional plasmids pCff2 and pCff3 present in ICMP 22053, were absent in all Australian *C. flaccumfaciens* pv*. flaccumfaciens* isolates in this study too. Interestingly, the above-mentioned 27 kb region of pCff1 and pCff2 and pCff3 are also absent in the Illumina assembly of ICMP 22053 released under the name CFBP 8820 (JAHEWR000000000; Osdaghi et al. (2022)), suggesting these may be an artefact of PacBio assembly.

We did not detect host specificity of MLLs as isolates obtained from different host crops occurred within the same MLLs. For example, MLL29 and MLL31 consisted of genotypes from black gram and cowpea. Moreover, isolates from different hosts were genetically very closely related (for example, DAR 56678 from cowpea and DAR 58696 from mungbean). However, only a limited number of isolates from hosts other than mungbean were included in this study and results need to be treated with caution.

Our phylogenetic analysis of single orthologous loci showed that CFBP 3417 and CFBP 3422, isolated from bean and originally submitted to NCBI as *C. flaccumfaciens* pv*. flaccumfaciens* (BioProject PRJNA731370), clade separately from other *C. flaccumfaciens* pv*. flaccumfaciens* isolates (Supplementary Fig. S7). Also, CFBP 8818, CFBP 8819, CFBP 8823, CFBP 8824, isolated from tomato and originally identified as *C. flaccumfaciens* pv*. flaccumfaciens,* did not form a monophyletic clade with *C. flaccumfaciens* pv*. flaccumfaciens*. This, together with the ANI values < 95%, confirmed the requirement to reclassify these into a separate genomospecies, in line with findings by Evseev et al. (2022a). Subsequent to our analyses, CFBP 8818, CFBP 8819, CFBP 8823, CFBP 8824 were described as *Curtobacterium aurantiacum* (Osdaghi et al. 2024).

The shift in published resistance ratings of mungbean cultivars Crystal and Celera from moderately resistant (MR) in 2008 (Gentry & Cumming 2008; Gentry 2010) to moderately susceptible (MS) in 2013 (Douglas & Cumming 2013, 2014; GRDC 2017) has not been investigated. While these changes could be interpreted as an erosion of resistance in these cultivars, our study found no evidence of generation or introduction of isolates with enhanced aggressiveness causing such shifts. The initial MR rating may have resulted from confusion of pseudoresistance with true genetic resistance. Pseudoresistance (or disease escape) in crops refers to a temporary appearance of resistance that is not due to inherent genetic resistance, and arises from external or environmental factors that suppress the activity of the pathogen or limit disease development (Pedersen 1960; Matre et al. 2022). While field screenings for tan spot resistance relied on spray inoculation, stem pinprocking has been reported to provide a better indication of resistance to *C. flaccumfaciens* pv*. flaccumfaciens*, which is a xylem-inhabiting pathogen (Diatloff & Imrie 2000). Spray inoculation simulates in-field infection of plants via splash dispersal of inoculum and can provide important information regarding varietal resistance to in-field leaf infection of mungbean plants. However, stem inoculation be used as the primary method of selection for tan spot resistance (Schuster & Coyne 1981; Diatloff & Imrie 2000). Our unpublished studies have also indicated leaf spraying may result in underestimation of disease severity.

It is possible that *C. flaccumfaciens* pv*. flaccumfaciens* isolates used for the initial resistance screening in the 2000s had lower aggressiveness resulting is higher resistance rating of some cultivars at the time. Population changes have occurred in another bacterial pathogen of mungbean in Australia, *Pseudomonas savastanoi* pv. *phaseolicola*, causal agent of halo blight (Noble et al. 2020), where the resistance of cultivars bred to be resistant to one isolate was overcome by the presence of a more aggressive isolate in the field. Unfortunately, *C. flaccumfaciens* pv*. flaccumfaciens* isolates used in the national mungbean improvement program in Australia in the 1990s and 2000s were not available to us to confirm their genotype or adaptive fitness. Also, the older mungbean genotypes (cv. Celera and Crystal) were not available, so, we could not further investigate this possibility.

To the best of our knowledge, the mungbean breeding program in Australia were making use of isolates CffUQ35, CffUQ36 and CffUQ37 for breeding efforts in 2018 and 2019, which our study showed belong to the same clonal lineage (Cluster 4), which had a low frequency (7%) in the Australian *C. flaccumfaciens pv. flaccumfaciens* population from mungbean fields. This lower frequency despite aggressiveness of these isolates could potentially be due to lower local adaptation of these isolates to Australian conditions or lower saprophytic fitness, making these unsuitable for field screening of mungbean cultivars for resistance to tan spot.

It should be noted that we tested aggressiveness of isolates separately, which is not necessarily indicative of competitive fitness of isolates. Development of species-specific and isolate-specific quantitative PCR assays will allow for more accurate and sensitive disease assessment though *in planta* quantification of the pathogen and enable investigation of competitive fitness of isolates from different genotypes when inoculated as mixtures, which is currently in progress.

This study emphasizes the importance of population studies to understand pathogen genetic and phenotypic diversity and support plant breeding programs. It is critical to understand the diversity of local populations and select dominant, aggressive, and locally adapted pathogen genotypes to use in phenotyping for resistance to achieve effective and durable resistance. Further research should focus on exploring the mechanisms underlying clonal expansion of *C. flaccumfaciens* pv*. flaccumfaciens* Cluster 1 in Australia, including investigation of the function of private alleles identified in this study and further competitive fitness assays.

## Materials and Methods

### Sampling strategy and bacterial isolations

In 2018 and 2019, sampling was conducted from 12 mungbean fields in Qld and northern NSW. Due to the lack of prior knowledge of the genetic diversity of *C. flaccumfaciens* pv*. flaccumfaciens* in Australia, two sampling strategies were adopted. The first strategy was applied in fields where disease severity was high (field 10 in Qld and field 42 in northern NSW), with hierarchically structured, intensive sample collection conducted along two transects. The transects were 50 m apart, with 15 sampling points along each transect at 15 m intervals. At all locations along the transects, only one symptomatic plant was sampled, except for the first location, where six symptomatic plants were sampled within a 5 m radius. This resulted in a total of 40 plants from each intensively sampled field. The second strategy was applied in fields where disease severity was low, with opportunistic sampling conducted by randomly collecting symptomatic leaves (sampling points at least 5 m apart from each other). A total of 200 symptomatic mungbean plants were collected from NSW and Qld.

Symptomatic leaves were surface sterilized by spraying with 70% ethanol and wiping with autoclaved paper towels. Leaf tissues were excised from the margin of lesions and cut in halves. One half was placed in a drop of water on a microscope slide and observed under a compound microscope and another half was placed in a microfuge tube containing 100 µl sterile water. Upon confirmation of bacterial oozing under the microscope, a sterile loop was used to streak the water suspension onto Nutrient Agar (NA) (Amyl Media, Victoria, Australia). For intensively sampled fields, multiple isolations (three to five) were conducted on subset of haphazardly selected leaves to obtain multiple isolates per lesion.

The identity of all isolates was assessed using species-specific PCR primers CF4/CF5 (Messenberg Guimaraés et al. 2001) as well as Sanger sequencing the 16S region of the nuclear ribosomal DNA using universal 16S primers 27F/1492R (Lane 1991; Turner et al. 1999). To explore the possible occurrence of colony types producing yellow, orange, pink and purple pigments (Agarkova et al. 2012) in Australia, we sequenced any yellow-, orange-, pink-, -red, and purple-pigmented colonies isolated from symptomatic mungbean leaves.

### *Curtobacterium flaccumfaciens* pv. *flaccumfaciens* isolates from Australian fields

A collection of 119 *C. flaccumfaciens* pv*. flaccumfaciens* isolates (Supplementary Table S1) was used for this whole genome sequencing study. This included 77 isolates collected in 2018 and 2019 from 12 fields as described above, 21 isolates collected from 2014 to 2017 from 14 fields, and 21 historical isolates collected from initial tan spot outbreaks in 1986 to 1987 from Qld and NSW (NSW Plant Pathology and Mycology Herbarium, NSW Department of Primary Industries [DAR Culture Collection, Orange, New South Wales]). Isolates originated from mungbean (*V. radiata*; *n* = 104), cowpea (*V. unguiculata*; *n* = 5), black gram (*V. mungo*; *n* = 3), soybean (*Glycine max; n* = 2), adzuki bean (*V. angularis*; *n* = 2), and common bean (*Phaseolus vulgaris*; *n* = 1).

### Genome sequencing

Genomic DNA was extracted using a PureLink® Genomic DNA Kit (Life Technologies, Victoria, Australia), according to manufacturer’s instructions. All DNA samples were assessed for integrity using agarose gel electrophoresis, quantified using a fluorometer (Qubit 3.0, dsDNA BR Kit, Thermo Fisher Scientific, Victoria, Australia), and stored at -20°C. Genomic DNA libraries were constructed from 200 ng of each DNA extract using Nextera^TM^ DNA Flex library prep kits and Nextera^TM^ DNA CD Indexes (Illumina, Singapore) for 130 DNA samples, which included 119 unique samples and an additional 11 replicated samples repeated within and between sequencing runs as positive controls. These positive controls belonged to DNA samples from the same isolates to assess sequencing errors and variation among sequencing runs. Illumina library preparations were conducted according to manufacturer’s instructions, with minor modifications. For PCR enrichment, the number of PCR cycles were reduced to four in order to reduce sequencing error and improve accuracy (Bruinsma et al. 2018). Libraries were pooled in batches of 48, 48, and 34 DNA samples, and quantified using a Qubit^TM^ dsDNA High Sensitivity kit (Thermo Fisher Scientific). Due to the high GC content of *Curtobacterium* spp. genomes (Klein et al. 2016; Vaghefi et al. 2021), the chemically denatured pooled library was exposed to heat denaturation (2 minutes incubation at 96°C followed by 5 minutes in an ice water bath) to increase the rate of cluster generation. Heat-denatured libraries were sequenced at molecular laboratories of the University of Southern Queensland, on an Illumina MiSeq platform using a 600-cycle paired-end V3 reagent kits (Illumina).

### Variant calling and filtering

Raw Illumina data for all DNA samples (*n* = 130) were filtered using BBduk program from BBTools version 1.0 (Bushnell 2018) to remove adapter sequences (*ktrim = r, k = 23, mink = 11, hdist = 1, minlen = 50, ftm = 5 tpe tbo*), low-quality nucleotides (*qtrim = r, trimq=10, minlen=50, maq=10 tpe*) and common sequence contaminants based on the NCBI Univec database (*k = 31, hdist = 1*).

The complete genome assembly of BRIP 70614 (NCBI genome accession no. GCA_017909455.1), an Australian *C. flaccumfaciens* pv*. flaccumfaciens* isolate from mungbean (Vaghefi et al. 2021), was selected for reference-based variant calling. Filtered sequences were aligned to the reference genome using BWA-MEM2 Galaxy version 2.2.1 (Li 2013) in Galaxy Australia portal (Jalili et al. 2020). Sorting and removal of PCR duplicates were conducted in SAMtools Galaxy version 1.15.1 (Li et al. 2009). Variant calling was conducted in Freebayes-parallel version 1.3.5 (Garrison & Marth 2012; Tange 2018) on the Linux cluster at the Cornell University BioHPC Computing Facilities (Ithaca, New York, USA). Variant-calling was conducted to retain variable sites only (*--ploidy* 1 *--min-alternate-fraction* 0.95 --*min-alternate-count* 10 *--min-mapping-quality* 20 --*min-base-quality* 30).

Preliminary filtering was conducted as follows. The average coverage per site (*--site-depth*) and missing data per individual (*--missing-ind*) were calculated using vcfTools version 0.1.16 (Danecek et al. 2011). VcfTools was further used to remove two genotypes that showed > 70% missing data (CffUQ19 and CffDA027). All remaining genotypes had < 9% missing data and were retained. SNPSift (Cingolani et al. 2012) from SNPEff version 4.3 (Cingolani et al. 2012) was used to filter out over-represented sites (threefold higher coverage than the average), because these could be the result of repetitive sequences and result in unreliable variants (McCann et al. 2017). To remove sequencing and genotyping errors, sites that showed different variant calls between replicated samples were removed using SNPSift. Finally, all replicated samples (*n* = 11) were removed using vcfTools, retaining 117 genotypes in the data set.

For population genomic analyses, the data set was further filtered to retain informative variants with a minimum allele frequency of 5% (*maf* = 0.05) and maximum missing data of 10%. This resulted in a total of 51,059 Single Nucleotide Polymorphisms (SNPs) across the 117 isolates, of which 50,960 were core SNPS (located on the core chromosome) and 7,174 were core indels (insertion/deletion). SNPs were annotated against gene models of the BRIP 70614 reference genome using SnpEff to identify variations with high (affecting splice-sites, stop and start codons), moderate (non-synonymous), low (synonymous coding/start/stop, start gained), or modifier (upstream, downstream, intergenic, UTR) impact. Of the 50,960 core SNPs, 316, 16,291, 30,571, and 3,782 were identified as high, moderate, low, and modifier impact SNPs, respectively. For population structure analyses, which require a set of selectively neutral markers, 30,035 core SNPs with low and modifier impact were used. Low impact SNPs do not to change protein function while modifier (intergenic) variants reside in non-coding regions of the genome; therefore, there is no evidence of impact (Cingolani et al. 2012) andboth groups are considered as potentially neutral SNPs. Population structure analyses were repeated a second time using all SNPs including predicted high and moderate impact to allow comparison with results obtained from the neutral loci. All other analyses were conducted using the entire set of SNPs after removing loci with missing data as described below.

### Genotypic diversity and spatial and temporal distribution of multi-locus genotypes (MLGs)

A critical step in population genomic analyses of clonal organisms is identification of clones, or unique multi-locus genotypes (MLGs) (Grünwald et al. 2016; Arnaud-Haond et al. 2007). MLG assignment using whole genome sequence data is challenging as sequencing errors, variant-calling errors, and missing data (Grünwald et al. 2016; Milgroom 2017) inflate the number of MLGs. In this study, sequencing and genotyping errors were minimised by including seven within-plate and four between-plate replicated DNA samples (Supplementary Table S8), which allowed us to remove unreliable sites that carried different alleles between replicated DNA samples. Moreover, all sites with missing data were removed using SNPfiltR version 1.0.1 (DeRaad 2022), resulting in 43,974 SNPs for analyses of genotypic diversity.

Accumulation of somatic mutations in clonally reproducing populations may result in unique MLGs that are differentiated based on only a handful of loci, resulting in overestimation of the number of MLGs, hence, inflated genotypic diversity (Milgroom 2017). Therefore, it is important to delineate clonal boundaries based on a defined genetic distance threshold and combine MLGs that have limited variation into multi-locus linages (MLLs) (Vaghefi et al. 2017; Kamvar et al. 2015; Milgroom 2017). We found that 20 unique MLGs (defined as isolates with the same SNP profile) in the Australian *C. flaccumfaciens* pv*. flaccumfaciens* population differed at only one to seven loci (Fig. S1). Thus, *cutoff_predictor* and *mlg.filter* functions in *poppr* version 2.9.3 (Kamvar et al. 2014) were used to collapse all MLGs with genetic distance smaller than the estimated threshold of 0.0001468889 (estimated by *cutoff_predictor*) into the same MLL (Kamvar et al. 2014, 2015). The *cutoff_predictor* function in *poppr* identifies the optimal genetic distance threshold where the number of observed MLGs plateaus to define distinct genotypes, while the *mlg.filter* function applies this threshold to group similar genotypes by filtering and clustering them based on their genetic distances. Thus, MLGs with one to seven SNP differences were collapsed into MLLs.

Nei’s measure of allelic diversity (H_e_) (Nei 1978), the number of multi-locus lineages (MLLs), expected number of MLLs at smallest samples size (eMLL), clonal fraction (defined as (N-number of MLLs)/N, where N is the total number of isolates), Simpson’s complement index of genotypic diversity corrected for sample size ((N/(N-1))*λ) (Simpson 1949), and recurrent MLLs (MLLs that occurred more than once) were obtained using *poppr*. For intensively sampled fields, a plot of total genotypic diversity against sample size was created by calculating Simson’s complement index (1-λ) of genetic diversity (Simpson 1949) for populations ranging in size from n to one. Populations from intensively sampled fields were clone-corrected to plant level in *poppr* and only unique MLGs were used for subsequent analyses to ensure repeated isolation of the same MLG did not skew the results.

Recurrent MLGs and MLLs (those shared among years, hosts, and locations) were identified in *poppr* using the *mlg.crosspop* function. MLL-specific (private) alleles, defined as alleles that are unique to one MLL, were identified in SNPSift using the entire SNP data set. The influence of geographic distance on genetic distance among MLGs was investigated by performing a Mantel test in R using the package *vegan* version 2.64 (Oksanen et al. 2024) based on the Pearson correlation coefficient with 999 permutations. The Mantel statistic, r, ranging from -1 to 1, measures the correlation between two dissimilarity or distance matrices. To investigate the correlation between genetic and geographic distance, while considering the influence of time, a partial Mantel test was conducted using the function *mantel.partial* in *vegan*.

### Population structure and temporal differentiation of *Curtobacterium flaccumfaciens* pv. *flaccumfaciens* in Australia

The existence of an underlying population structure without *a priori* assumption of populations was tested using the model-based Bayesian clustering method implemented in STRUCTURE version 2.3.4 (Pritchard et al. 2000), which is not sensitive to clonality or linkage of loci. Assignment of isolates to clusters was inferred for one to 10 genetic groups (*K*), and each *K* value model was replicated 10 times (iterations) with a burn-in period of 100,000 steps, followed by 1,000,000 Markov chain Monte Carlo steps. The optimal *K* value was chosen by computing Δ*K* using the method proposed by (Evanno et al. 2005) in Microsoft Excel. The 10 replicated runs for the optimal *K* value were combined and a single graphical output was generated using CLUMPAK version 1.1 (Kopelman et al. 2015).

We performed Fisher’s Exact test of independence to investigate statistically significant changes in the frequency of identified clusters between the historical (1986 and 1987) and contemporary (2014 to 2019) populations in SPSS version 21. Adjusted standardized residuals calculated in SPSS as part of a post-hoc analysis were used to examine statistically significant differences in the frequency of isolates between populations.

A Discriminant Analysis of Principal Components (DAPC) was conducted using *adegenet* version 2.1.10 (Jombart 2008), with the optimal number of clusters determined using the function *find.clusters* in *poppr*. DAPC was then used to assign individuals into clusters, retaining the number of principal components encompassing 97% of the cumulative variance.

### Phylodynamic analysis

Bayesian phylodynamic analysis was conducted in Bayesian Evolutionary Analyses of Sampled Trees (BEAST) version 2.7.7 (Bouckaert et al. 2019) to reconstruct evolutionary dynamics of the Australian *C. flaccumfaciens* pv*. flaccumfaciens* population over time, providing insights into demographic changes and the temporal spread of the pathogen. For this, the clone-corrected core SNP alignment was used with tip dates defined as the year of collection. A HKY nucleotide substitution model with empirical base frequencies and gamma distribution of site-specific rate heterogeneity was used. The analysis was repeated under a strict molecular clock using both Coalescent Constant Size and Exponential Growth tree priors to investigate the potential impact of priors on phylogenies Bayesian Markov chain Monte Carlo analyses were run for 100 million steps and samples were drawn every 10,000 MCMC. The MCMC convergence was assessed based on a minimum effective sample size of 150 for parameter estimates using Tracer version 1.7.2. A maximum clade credibility tree was generated using TreeAnnotator version 2.7.7 after a burn-in of 25%. Phylogenies were visualized in FigTree version 1.4.4 (http://tree.bio.ed.ac.uk/software/figtree/).

### Outlier analyses for detection of loci potentially under selection

Outlier loci are genetic regions that show unusual patterns of variation, often indicating they are under selective pressure or influenced by environmental factors. In order to identify SNPs potentially under selection from 1986, F_st_ outlier analysis was conducted using pairwise comparison of the historical (1986 and 1987) and contemporary (2014 to 2019) populations to detect loci that show unusual patterns of variation between the two populations. F_st_ is a measure of genetic differentiation between populations, quantifying the proportion of genetic variation that is distributed among populations rather than within them, with values ranging from 0 (no differentiation) to 1 (complete differentiation). SNP-by-SNP F_st_ analysis, which detects outlier loci by comparing genetic differentiation at individual SNPs between populations, was conducted using a sliding window approach in vcfTools (*--fst-window-size* 50,000 *--fst-window-step* 50,000). F_st_ values for all windows were subsequently plotted across the chromosome to generate a Manhattan plot in R using the library *ggplot2* (Wickham 2016) (Supplementary Figure S9). The top 1% of SNPs (greater than the quantile value of 99%) were classified as outliers to reduce the probability of false positives.

BayeScan version 2.1 (Foll & Gaggiotti 2008) was used as a second approach to detect outlier loci potentially under selection since 1986. This method identifies outliers based on allele frequencies, taking into account the underlying population structure. BayeScan was run with 20 pilot runs of 5,000 iterations each followed by a main run of 5,000 iterations, a thinning interval of 10, with a burn in of 50,000 and an FDR of 1%, 5% or 10% to identify outliers, which were further characterized for potential function.

### Australian *Curtobacterium flaccumfaciens* pv. *flaccumfaciens* population within a global context

Published genome assemblies of 19 *C. flaccumfaciens* pv*. flaccumfaciens* (taxon 138532) isolates available in December 2023 were retrieved from the NCBI database to be used as reference *C. flaccumfaciens* pv*. flaccumfaciens* genomes (Supplementary Table S6). Alignment of core genes was carried out as described by (Colombi et al. 2023). Briefly, all genomes were annotated with Prokka version 1.14.6 (Seemann 2014). Proteinortho version 6.3.0 was used to identify single-copy orthologues and their nucleotide sequences were employed for codon-aware alignment with PRANK version 170427 (Löytynoja & Russell 2014). Core gene alignments were concatenated and stripped of gaps with Goalign version 0.3.7 (https://github.com/evolbioinfo/goalign).

First, a Neighbor-Net phylogenetic network was constructed using SplitsTree version 4.18.3 (Huson & Bryant 2006) with 1,000 bootstrap replicates, which allows for recombination between nodes, and thus provides a more accurate representation of recombining data sets. Also, Maximum-likelihood phylogeny reconstruction was then performed with RAxML version 7.2.8 (Stamatakis 2014). ClonalFrameML version 1.178 (Didelot & Wilson 2015) was employed to identify polymorphisms introduced via homologous recombination using the gap-free core genes alignment and initial maximum-likelihood phylogeny as input. Alignment regions identified as likely to have been introduced via recombination were removed from the alignment and tree building was repeated using the non-recombinant alignment with RAxML (*-f a -p $RANDOM -x $RANDOM -# 100 m GTRCATX -T 16*). Trees were visualized using the R package ggtree version 3.4.4 (Yu et al. 2017). FastANI version 1.34 was used to calculate average pairwise nucleotide identity between the *C. flaccumfaciens* pv*. flaccumfaciens* isolates and the *C. flaccumfaciens* pv*. flaccumfaciens* type isolate LMG 3645. Carlos7, having an ANI < 95% in comparison to the *C. flaccumfaciens* pv. *flaccumfaciens* type strain LMG 3645, has been suggested to belong to a different species (Evseev et al. 2022a), and was used to root the *C. flaccumfaciens* pv*. flaccumfaciens* phylogeny.

### Plasmid detection and phylogeny

*Curtobacterium flaccumfaciens* pv*. flaccumfaciens* genomes annotated by Prokka were used to identify the presence of mobile genetic elements as follows. Initially, conjugation systems were searched in the Prokka-annotated genomes with CONJscan version 1.0.2 (Cury et al. 2020) using default parameters. After CONJscan failed to recognise any known conjugation systems, we looked for relaxase gene sequences to identify putative horizontally transmissible elements as described in Colombi et al. (2021). Relaxase gene sequences were searched with hmmscan in HMMER version 1.2.1 (Eddy 2011) using available hidden Markov model (HMM) protein profiles (MOB database) (Abby & Rocha 2017) defined for distinct MOB-protein families (Francia et al. 2004) with a bit score threshold (-T) of 33.

To identify known plasmid sequences, *C. flaccumfaciens* pv*. flaccumfaciens* genome sequences were interrogated with BLASTn version 2.7.1 (Altschul et al. 1990) with default parameters using pCff119 (CP074440.1), pCff2 (CP045289.2) and pCff3 (CP045290.2) as queries. Retrieved sequences were aligned to pCff119 using MAFFT version 7.520 (Katoh 2002) with default parameters. The method used to infer the phylogeny of the plasmids was the same as that used for the *C. flaccumfaciens* pv*. flaccumfaciens* phylogeny.

The plasmid SNPs dataset (99 SNPs detected on the plasmid in the Australian collection) was used to construct a plasmid haplotypic network. Plasmid haplotypes in the Australian isolates and their frequencies across different years were depicted using the software POPART version 1.7 (Leigh & Bryant 2015).

### Pathogenicity and aggressiveness assays

Three isolates lacking the linear plasmid (DAR 58700, Cff089, and BRIP 70624) and three isolates carrying the linear plasmid (BRIP 70606, BRIP 70623, and BRIP 70614) were selected for pathogenicity tests on black gram (*Vigna mungo* cv. Onyx-AU), cowpea (*Vigna unguiculata* cv. Red Caloona), and mungbean (*Vigna radiata* cv. Jade-AU). A completely randomised trial included three replicates per isolate. Plant growth, inoculum preparation and inoculation were conducted according to Vaghefi (2021). Plants were assessed for symptom development on a daily basis. The pathogen was reisolated from symptomatic tissue and its identity as *C. flaccumfaciens* pv*. flaccumfaciens* was confirmed through 16S sequencing.

To test the hypothesis of variation in aggressiveness of isolates from different clusters or carrying different plasmid haplotypes, 13 isolates (two to three from each cluster) were selected to test for pathogenicity and aggressiveness on mungbean cv. Jade-AU (Table 1).

The experiment followed a completely randomised design in controlled environment rooms under 16/8 hr light/darkness photoperiod at 27/18°C (day/night temperature) and 70% humidity. Inoculum preparation and inoculation were conducted according to Vaghefi (2021). The experiment was repeated two times including four and five internal replications in trial 1 and 2, respectively. Disease severity was assessed at 25 days after inoculation by estimating the average symptomatic leaf area (percent diseased area) of the first two trifoliates of each plant using the software Leaf Doctor (Pethybridge & Nelson 2015) as described by Vaghefi (2021). The pathogen was reisolated from symptomatic tissue and its identity as *C. flaccumfaciens* pv*. flaccumfaciens* was confirmed through 16S sequencing. The statistical analyses and significance of disease severity caused by different *C. flaccumfaciens* pv. *flaccumfaciens* isolates was tested using the non-parametric Kruskal Wallis test in SPSS version 21. Pairwise comparisons were conducted using Mann-Whitney U tests with significance values adjusted by the Bonferroni correction for multiple tests in SPSS. Both trials were included in the one

## Acknowledgements

We would like to thank Ms. Encarnacion Adorada for excellent technical support. This project was funded by the Grains Research and Development Corporation Project DAQ0018 and Queensland Government Department of Agriculture and Fisheries Broadacre Cropping Initiative Project 36. We thank the four anonymous reviewers for insightful feedback that improved the manuscript significantly.

## Authors’ contributions

The work presented resulted from collaboration between all authors and each author has participated sufficiently in the work to take responsibility for the content. AHS and NV conceived and designed the project. AY, DLA, LAK, and NV contributed to sample collection and isolation. DLA performed DNA extractions and quality control. AY and NV conducted whole genome sequencing of isolates. NV and NLK carried out the pathogenicity assays. AHS, NV and EC conducted the data analyses and drafted the manuscript. All co-authors provided revisions of the manuscript.

## Data Availability

Data sets are publicly available on Zenodo DOI: 10.5281/zenodo.10440730

## Notes

https://zenodo.org/records/10440730

### Competing Interest Statement

The authors have declared no competing interest.

## References

Abby SS et al. 2016. Identification of protein secretion systems in bacterial genomes. Sci. Rep. 6:23080. doi: 10.1038/srep23080.

Abby SS, Rocha EP. 2017. Identification of protein secretion systems in bacterial genomes using MacSyFinder. Bact. Protein Secret. Syst. Methods Protoc. 1–21.

Agarkova IV, Lambrecht PA, Vidaver AK, Harveson RM. 2012. Genetic diversity among *Curtobacterium flaccumfaciens* pv. *flaccumfaciens* populations in the American High Plains. Can. J. Microbiol. 58:788–801. doi: 10.1139/w2012-052.

Altschul SF, Gish W, Miller W, Myers EW, Lipman DJ. 1990. Basic local alignment search tool. J. Mol. Biol. 215:403–410. doi: 10.1016/S0022-2836(05)80360-2.

Arnaud-Haond S, Duarte CM, Alberto F, Serrão EA. 2007. Standardizing methods to address clonality in population studies. Mol. Ecol. 16:5115–5139. doi: 10.1111/j.1365-294X.2007.03535.x.

Bouckaert R et al. 2019. BEAST 2.5: An advanced software platform for Bayesian evolutionary analysis Pertea, M, editor. PLOS Comput. Biol. 15:e1006650. doi: 10.1371/journal.pcbi.1006650.

Bruinsma S et al. 2018. Bead-linked transposomes enable a normalization-free workflow for NGS library preparation. BMC Genomics. 19:722. doi: 10.1186/s12864-018-5096-9.

Bushnell B. 2018. BBTools: a suite of fast, multithreaded bioinformatics tools designed for analysis of DNA and RNA sequence data. sourceforge.net/projects/bbmap/.

Castillo AI et al. 2021. Allopatric Plant Pathogen Population Divergence following Disease Emergence Cann, I, editor. Appl. Environ. Microbiol. 87:e02095–20. doi: 10.1128/AEM.02095-20.

Chen G et al. 2021. Complete Genome Sequencing Provides Novel Insight Into the Virulence Repertories and Phylogenetic Position of Dry Beans Pathogen *Curtobacterium flaccumfaciens* pv. *flaccumfaciens*. Phytopathology®. 111:268–280. doi: 10.1094/PHYTO-06-20-0243-R.

Cingolani P, Platts A, et al. 2012. A program for annotating and predicting the effects of single nucleotide polymorphisms, SnpEff: SNPs in the genome of Drosophila melanogaster strain w ^1118^ ; iso-2; iso-3. Fly (Austin). 6:80–92. doi: 10.4161/fly.19695.

Cingolani P, Patel VM, et al. 2012. Using Drosophila melanogaster as a Model for Genotoxic Chemical Mutational Studies with a New Program, SnpSift. Front. Genet. 3. doi: 10.3389/fgene.2012.00035.

Collins MD, Jones D. 1983. Reclassification of Corynebacterium flaccumfaciens, Corynebacterium betae, Corynebacterium oortii and Corynebacterium poinsettiae in the genus Curtobacterium, as Curtobacterium flaccumfaciens comb. nov. Microbiology. 129:3545–3548. doi: 10.1099/00221287-129-11-3545.

Colombi E et al. 2021. Comparative analysis of integrative and conjugative mobile genetic elements in the genus Mesorhizobium. Microb. Genomics. 7. doi: 10.1099/mgen.0.000657.

Colombi E et al. 2023. Population genomics of Australian indigenous Mesorhizobium reveals diverse nonsymbiotic genospecies capable of nitrogen-fixing symbioses following horizontal gene transfer. Microb. Genomics. 9.

Cury J, Abby SS, Doppelt-Azeroual O, Néron B, Rocha EPC. 2020. Identifying Conjugative Plasmids and Integrative Conjugative Elements with CONJscan. In: Horizontal Gene Transfer. De La Cruz, F, editor. Methods in Molecular Biology Vol. 2075 Springer US: New York, NY pp. 265–283. doi: 10.1007/978-1-4939-9877-7_19.

Danecek P et al. 2011. The variant call format and VCFtools. Bioinformatics. 27:2156–2158. doi: 10.1093/bioinformatics/btr330.

DeRaad DA. 2022. SNPFILTR : An R package for interactive and reproducible SNP filtering. Mol. Ecol. Resour. 22:2443–2453. doi: 10.1111/1755-0998.13618.

Diatloff A, Imrie BC. 2000. [No title found]. Australas. Plant Pathol. 29:24. doi: 10.1071/AP00004.

Didelot X, Wilson DJ. 2015. ClonalFrameML: efficient inference of recombination in whole bacterial genomes. PLoS Comput. Biol. 11:e1004041.

Douglas C, Cumming G. 2013. Australian Mungbean Association. 2013. Pulse Variety Management Package, Jade-AU, Large seeded shiny green mungbean. http://www.mungbean.org.au/assets/2013_vmp_jade-au.pdf.

Douglas C, Cumming G. 2014. Celera II-AU, small seeded shiney green mungbean, high grain quality improved halo blight resistance.

Eddy S. 2011. Accelerated profile HMM searches. PLOS Computational Biology 7.

Evanno G, Regnaut S, Goudet J. 2005. Detecting the number of clusters of individuals using the software STRUCTURE : a simulation study. Mol. Ecol. 14:2611–2620. doi: 10.1111/j.1365-294X.2005.02553.x.

Evseev P et al. 2022a. Curtobacterium spp. and Curtobacterium flaccumfaciens: Phylogeny, Genomics-Based Taxonomy, Pathogenicity, and Diagnostics. Curr. Issues Mol. Biol. 44:889–927. doi: 10.3390/cimb44020060.

Evseev P et al. 2022b. Curtobacterium spp. and Curtobacterium flaccumfaciens: Phylogeny, Genomics-Based Taxonomy, Pathogenicity, and Diagnostics. Curr. Issues Mol. Biol. 44:889–927. doi: 10.3390/cimb44020060.

Foll M, Gaggiotti O. 2008. A Genome-Scan Method to Identify Selected Loci Appropriate for Both Dominant and Codominant Markers: A Bayesian Perspective. Genetics. 180:977–993. doi: 10.1534/genetics.108.092221.

Francia, et al. 2004. A classification scheme for mobilization regions of bacterial plasmids. FEMS Microbiol. Rev. 28:79–100. doi: 10.1016/j.femsre.2003.09.001.

Garrison E, Marth G. 2012. Haplotype-based variant detection from short-read sequencing. doi: 10.48550/ARXIV.1207.3907.

Gentry J. 2010. Mungbean management guide 2nd edition.

Gentry J, Cumming G. 2008. Pulse Variety Management Package, Crystal, Large-seeded Bright Green Mungbean. http://www.mungbean.org.au/assets/2008_vmp_crystal.pdf.

Gonçalves RM et al. 2019. Genetic diversity of Curtobacterium flaccumfaciens revealed by multilocus sequence analysis. Eur. J. Plant Pathol. 154:189–202. doi: 10.1007/s10658-018-01648-0.

Goodwin SB, Cohen BA, Fry WE. 1994. Panglobal distribution of a single clonal lineage of the Irish potato famine fungus. Proc. Natl. Acad. Sci. 91:11591– 11595. doi: 10.1073/pnas.91.24.11591.

GRDC. 2017. Grownotes: Mungbeans. Grains Research and Development Corporation https://grdc.com.au/_data/assets/pdf_file/0014/315311/GRDC-GrowNotes-Mungbeans-Northern.pdf.

Grünwald NJ, McDonald BA, Milgroom MG. 2016. Population Genomics of Fungal and Oomycete Pathogens. Annu. Rev. Phytopathol. 54:323–346. doi: 10.1146/annurev-phyto-080614-115913.

Harveson RM, Schwartz HF. 2007. Bacterial Diseases of Dry Edible Beans in the Central High Plains. Plant Health Prog. 8:35. doi: 10.1094/PHP-2007-0125-01-DG.

Hedges F. 1922. A Bacterial Wilt of the Bean Caused by *Bacterium flaccumfaciens* Nov. Sp. Science. 55:433–434. doi: 10.1126/science.55.1425.433.

Helekal D, Ledda A, Volz E, Wyllie D, Didelot X. 2022. Bayesian Inference of Clonal Expansions in a Dated Phylogeny Ho, S, editor. Syst. Biol. 71:1073–1087. doi: 10.1093/sysbio/syab095.

Huson DH, Bryant D. 2006. Application of Phylogenetic Networks in Evolutionary Studies. Mol. Biol. Evol. 23:254–267. doi: 10.1093/molbev/msj030.

Jain C, Rodriguez-R LM, Phillippy AM, Konstantinidis KT, Aluru S. 2018. High throughput ANI analysis of 90K prokaryotic genomes reveals clear species boundaries. Nat. Commun. 9:5114.

Jalili V et al. 2020. The Galaxy platform for accessible, reproducible and collaborative biomedical analyses: 2020 update. Nucleic Acids Res. 48:W395– W402. doi: 10.1093/nar/gkaa434.

Jombart T. 2008. *adegenet* : a R package for the multivariate analysis of genetic markers. Bioinformatics. 24:1403–1405. doi: 10.1093/bioinformatics/btn129.

Kamvar ZN, Brooks JC, GrÃ¼nwald NJ. 2015. Novel R tools for analysis of genome-wide population genetic data with emphasis on clonality. Front. Genet. 6. doi: 10.3389/fgene.2015.00208.

Kamvar ZN, Tabima JF, Grünwald NJ. 2014. *Poppr* : an R package for genetic analysis of populations with clonal, partially clonal, and/or sexual reproduction. PeerJ. 2:e281. doi: 10.7717/peerj.281.

Katoh K. 2002. MAFFT: a novel method for rapid multiple sequence alignment based on fast Fourier transform. Nucleic Acids Res. 30:3059–3066. doi: 10.1093/nar/gkf436.

Kelly LA et al. 2021. One Crop Disease, How Many Pathogens? Podosphaera xanthii and Erysiphe vignae sp. nov. Identified as the Two Species that Cause Powdery Mildew of Mungbean (Vigna radiata) and Black Gram (V. mungo) in Australia. Phytopathology®. 111:1193–1206. doi: 10.1094/PHYTO-12-20-0554-R.

Klein BA et al. 2016. Draft Genome Sequence of *Curtobacterium* sp. Strain UCD-KPL2560 (Phylum *Actinobacteria*). Genome Announc. 4:e01040–16. doi: 10.1128/genomeA.01040-16.

Kopelman NM, Mayzel J, Jakobsson M, Rosenberg NA, Mayrose I. 2015. CLUMPAK : a program for identifying clustering modes and packaging population structure inferences across *K*. Mol. Ecol. Resour. 15:1179–1191. doi: 10.1111/1755-0998.12387.

Lane DJ. 1991. 16S/23S rRNA Sequencing. In: Nucleic acid techniques in bacterial systematics. In: Nucleic acid techniques in bacterial systematics. Stackebrandt, E & Goodfellow, M, editors. John Wiley and Sons: New York pp. 115–175.

Leigh JW, Bryant D. 2015. POPART : full-feature software for haplotype network construction Nakagawa, S, editor. Methods Ecol. Evol. 6:1110–1116. doi: 10.1111/2041-210X.12410.

Li H. 2013. Aligning sequence reads, clone sequences and assembly contigs with BWA-MEM. doi: 10.48550/ARXIV.1303.3997.

Li H et al. 2009. The Sequence Alignment/Map format and SAMtools. Bioinformatics. 25:2078–2079. doi: 10.1093/bioinformatics/btp352.

Li Y et al. 2012. Population Dynamics of *Phytophthora infestans* in the Netherlands Reveals Expansion and Spread of Dominant Clonal Lineages and Virulence in Sexual Offspring. G3 GenesGenomesGenetics. 2:1529–1540. doi: 10.1534/g3.112.004150.

Löytynoja A, Russell D. 2014. Multiple sequence alignment methods. Phylogeny-Aware Alignment PRANK.

Mable BK. 2019. Conservation of adaptive potential and functional diversity: integrating old and new approaches. Conserv. Genet. 20:89–100. doi: 10.1007/s10592-018-1129-9.

Matre YB, Zanwar P, Sonkamble MM. 2022. Host Plant Resistance Types and Its Mechanism.

McCann HC et al. 2017. Origin and Evolution of the Kiwifruit Canker Pandemic. Genome Biol. Evol. 9:932–944. doi: 10.1093/gbe/evx055.

McDonald BA, Stukenbrock EH. 2016. Rapid emergence of pathogens in agro-ecosystems: global threats to agricultural sustainability and food security. Philos. Trans. R. Soc. B Biol. Sci. 371:20160026. doi: 10.1098/rstb.2016.0026.

Messenberg Guimaraés P, Palmano S, Smith JJ, Grossi De Sá MF, Saddler GS. 2001. Development of a PCR test for the detection of Curtobacterium flaccumfaciens pv. flaccumfaciens. Antonie Van Leeuwenhoek. 80:1–10. doi: 10.1023/A:1012077425747.

Milgroom MG. 2017. Population Biology of Plant Pathogens: Genetics, Ecology, and Evolution. The American Phytopathological Society doi: 10.1094/9780890544525.

Nei M. 1978. ESTIMATION OF AVERAGE HETEROZYGOSITY AND GENETIC DISTANCE FROM A SMALL NUMBER OF INDIVIDUALS. Genetics. 89:583–590. doi: 10.1093/genetics/89.3.583.

Noble TJ et al. 2020. Characterisation of the Pseudomonas savastanoi pv. phaseolicola population found in Eastern Australia associated with halo blight disease in Vigna radiata. Australas. Plant Pathol. 49:515–524. doi: 10.1007/s13313-020-00722-8.

Oksanen J et al. 2024. Vegan: Community Ecology Package; R Package Version 2.6-8. https://github.com/vegandevs/vegan.

O’Leary ML, Gilbertson RL. 2020. Complete Genome Sequence Resource of a Strain of *Curtobacterium flaccumfaciens* pv. *flaccumfaciens*, the Causal Agent of Bacterial Wilt of Common Bean, from Turkey. Phytopathology®. 110:2010– 2013. doi: 10.1094/PHYTO-04-20-0131-A.

Osdaghi E et al. 2018a. Epiphytic *Curtobacterium flaccumfaciens* strains isolated from symptomless solanaceous vegetables are pathogenic on leguminous but not on solanaceous plants. Plant Pathol. 67:388–398. doi: 10.1111/ppa.12730.

Osdaghi E et al. 2018b. Epiphytic *Curtobacterium flaccumfaciens* strains isolated from symptomless solanaceous vegetables are pathogenic on leguminous but not on solanaceous plants. Plant Pathol. 67:388–398. doi: 10.1111/ppa.12730.

Osdaghi E et al. 2024. Multiphasic investigations imply transfer of orange-/red-pigmented strains of the bean pathogen Curtobacterium flaccumfaciens pv. flaccumfaciens to a new species as C. aurantiacum sp. nov., elevation of the poinsettia pathogen C. flaccumfaciens pv. poinsettiae to the species level as C. Poinsettiae sp. nov., and synonymy of C. albidum with C. citreum. Syst. Appl. Microbiol. 47:126489. doi: 10.1016/j.syapm.2024.126489.

Osdaghi E et al. 2016. Occurrence and characterization of a new red-pigmented variant of Curtobacterium flaccumfaciens, the causal agent of bacterial wilt of edible dry beans in Iran. Eur. J. Plant Pathol. 146:129–145. doi: 10.1007/s10658-016-0900-3.

Osdaghi E et al. 2022. Whole Genome Resources of 17 *Curtobacterium flaccumfaciens* Strains Including Pathotypes of *C. flaccumfaciens* pv. *betae*, C. flaccumfaciens pv. oortii, and C. flaccumfaciens pv. poinsettiae. Mol. Plant-Microbe Interactions®. 35:352–356. doi: 10.1094/MPMI-11-21-0282-A.

Osdaghi E, Lak MR. 2015. Occurrence of a New Orange Variant of *Curtobacterium flaccumfaciens* pv *. flaccumfaciens*, Causing Common Bean Wilt in Iran. J. Phytopathol. 163:867–871. doi: 10.1111/jph.12322.

Osdaghi E, Taghavi SM, Fazliarab A, Elahifard E, Lamichhane JR. 2015. Characterization, geographic distribution and host range of Curtobacterium flaccumfaciens: An emerging bacterial pathogen in Iran. Crop Prot. 78:185–192. doi: 10.1016/j.cropro.2015.09.015.

Osdaghi E, Young AJ, Harveson RM. 2020. Bacterial wilt of dry beans caused by *Curtobacterium flaccumfaciens* pv. *flaccumfaciens* : A new threat from an old enemy. Mol. Plant Pathol. 21:605–621. doi: 10.1111/mpp.12926.

Pedersen PN. 1960. Methods of Testing the Pseudo-resistance of Barley to Infection by Loose Smut, Ustilago nuda (Jens.) Rostr. Acta Agric. Scand. 10:312–332. doi: 10.1080/00015126009433045.

Pethybridge SJ, Nelson SC. 2015. Leaf Doctor: A New Portable Application for Quantifying Plant Disease Severity. Plant Dis. 99:1310–1316. doi: 10.1094/PDIS-03-15-0319-RE.

Pilik R et al. 2023. First Report of *Curtobacterium flaccumfaciens* pv. *flaccumfaciens* Causing Bacterial Wilt and Blight on Sunflower in Russia. Plant Dis. 107:1621. doi: 10.1094/PDIS-05-22-1203-PDN.

Pritchard JK, Stephens M, Donnelly P. 2000. Inference of Population Structure Using Multilocus Genotype Data. Genetics. 155:945–959. doi: 10.1093/genetics/155.2.945.

Savory EA et al. 2017. Evolutionary transitions between beneficial and phytopathogenic Rhodococcus challenge disease management. Elife. 6:e30925.

Schuster ML, Coyne DP. 1981. Biology, Epidemiology, Genetics and Breeding for Resistance to Bacterial Pathogens of Phaseolus vulgaris L. In: Horticultural Reviews. Janick, J, editor. Wiley pp. 28–58. doi: 10.1002/9781118060766.ch2.

Seemann T. 2014. Prokka: rapid prokaryotic genome annotation. Bioinformatics. 30:2068–2069.

Simpson EH. 1949. Measurement of Diversity. Nature. 163:688–688. doi: 10.1038/163688a0.

Stamatakis A. 2014. RAxML version 8: a tool for phylogenetic analysis and post-analysis of large phylogenies. Bioinformatics. 30:1312–1313.

Stukenbrock EH, McDonald BA. 2008. The Origins of Plant Pathogens in Agro-Ecosystems. Annu. Rev. Phytopathol. 46:75–100. doi: 10.1146/annurev.phyto.010708.154114.

Tange O. 2018. *Gnu Parallel* 2018. Zenodo doi: 10.5281/ZENODO.1146014.

Tarakanov RI et al. 2023. First Report of *Curtobacterium flaccumfaciens* pv. *flaccumfaciens* Causing Bacterial Tan Spot of Soybean in Russia. Plant Dis. 107:2211. doi: 10.1094/PDIS-08-22-1778-PDN.

Tegli S, Sereni A, Surico G. 2002. PCR-based assay for the detection of Curtobacterium flaccumfaciens pv. flaccumfaciens in bean seeds. Lett. Appl. Microbiol. 35:331–337. doi: 10.1046/j.1472-765X.2002.01187.x.

Turner S, Pryer KM, Miao VPW, Palmer JD. 1999. Investigating Deep Phylogenetic Relationships among Cyanobacteria and Plastids by Small Subunit rRNA Sequence Analysis. J. Eukaryot. Microbiol. 46:327–338. doi: 10.1111/j.1550-7408.1999.tb04612.x.

Vaghefi N et al. 2017. Global genotype flow in Cercospora beticola populations confirmed through genotyping-by-sequencing Chiang, T-Y, editor. PLOS ONE. 12:e0186488. doi: 10.1371/journal.pone.0186488.

Vaghefi N. 2021. Inoculating mungbean with Curtobacterium flaccumfaciens pv. flaccumfaciens (Cff) for tan spot disease assessment v1. doi: 10.17504/protocols.io.bym2pu8e.

Vaghefi N et al. 2015. Rapid Changes in the Genetic Composition of *Stagonosporopsis tanaceti* Population in Australian Pyrethrum Fields. Phytopathology®. 105:358–369. doi: 10.1094/PHYTO-08-14-0212-R.

Vaghefi N et al. 2019. What can we learn from population genomics studies of Curtobacterium flaccumfaciens pv. flaccumfaciens, the cause of tan spot on mungbean? In: Australasian Plant Pathology Society: Melbourne. http://era.daf.qld.gov.au/id/eprint/7161/.

Vaghefi N et al. 2021. Whole-Genome Data from *Curtobacterium flaccumfaciens* pv. *flaccumfaciens* Strains Associated with Tan Spot of Mungbean and Soybean Reveal Diverse Plasmid Profiles. Mol. Plant-Microbe Interactions®. 34:1216– 1222. doi: 10.1094/MPMI-05-21-0116-A.

Wickham H. 2016. ggplot2: Elegant Graphics for Data Analysis.

Wood B, Easdown W. 1990. A New Bacterial Disease of Mung Bean and Cowpea for Australia. Australas. Plant Pathol. 19:16. doi: 10.1071/APP9900016.

Yu G, Smith DK, Zhu H, Guan Y, Lam TT-Y. 2017. ggtree: an R package for visualization and annotation of phylogenetic trees with their covariates and other associated data. Methods Ecol. Evol. 8:28–36.

