## Supplementary_figures_and_tables for "Clonal expansion from standing genetic variation underpins the evolution of an emerging plant pathogen in Australia"

Adam H. Sparks[1,2,*], Dante L. Adorada[1], Elena Colombi[3,4,5], Lisa A. Kelly[1,6], Anthony Young[7], Noel L. Knight[1], and Niloofar Vaghefi[1,3**]

[1]Centre for Crop Health, University of Southern Queensland, Queensland, Australia
[2]Department of Primary Industries and Regional Development, Western Australia, Australia
[3]School of Agriculture, Food and Ecosystem Sciences, Faculty of Science, University of Melbourne, Victoria, Australia
[4]School of Natural Sciences, Macquarie University, Sydney, NSW, Australia
[5]ARC Centre of Excellence in Synthetic Biology, Sydney, NSW, Australia
[6]Department of Primary Industries, Queensland, Australia
[7]School of Agriculture and Food Sustainability, The University of Queensland, Queensland, Australia
*Current Address: Curtin Biometry and Agriculture Data Analysis, Curtin University, Bentley, Western Australian, Australia

### Contents

Supplementary Figures 2
Supplementary Tables 13

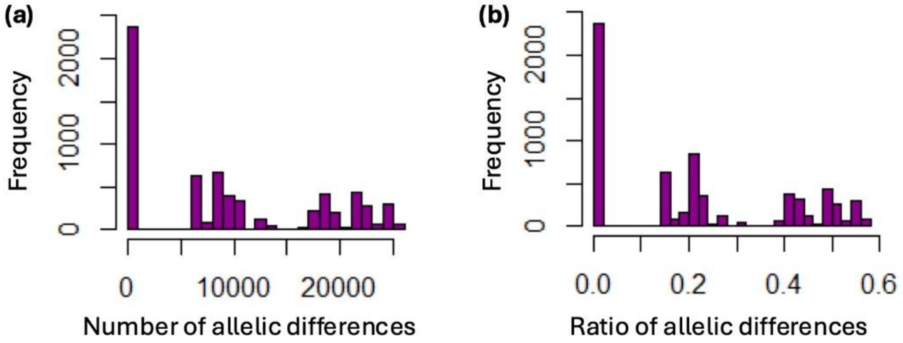


**Fig. S1.** Histograms of genetic distances between *Curtobacterium flaccumfaciens* pv. *flaccumfaciens* isolates collected from 1986 to 2019 in Australia, estimated via *diss.dist* function in *poppr* version 2.9.3 (Kamvar et al. 2014, 2015). **(a)** Number of single nucleotide polymorphism (SNP) differences for all pairwise comparisons of isolates **(b)** Ratio of the number of observed differences by the number of possible differences.

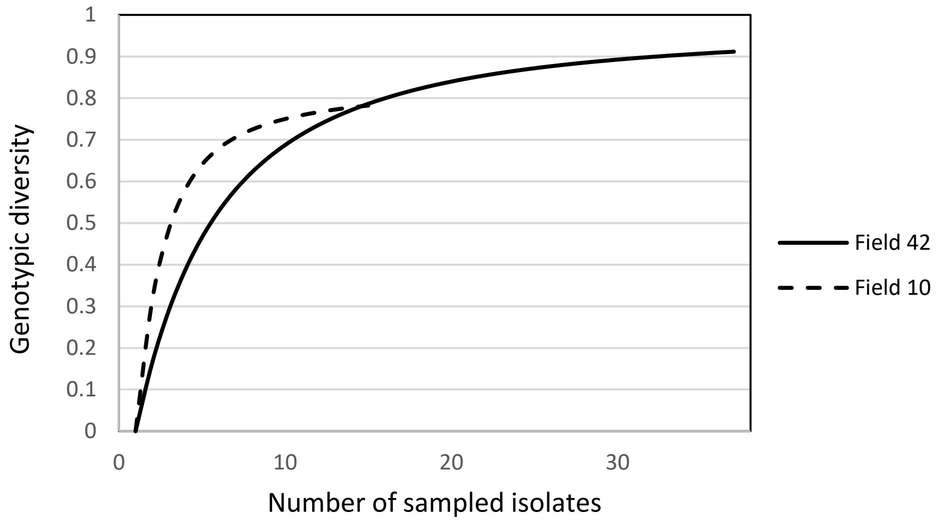


**Fig. S2.** Genotypic diversity estimated as Simpson's complement index of genotypic diversity (1-λ) (derived from Simpson’s (1949) λ) of *Curtobacterium flaccumfaciens* pv. *flaccumfaciens* populations by sample size in two intensively sampled mungbean fields in Queensland (Field 10) and New South Wales (Field 42), Australia, calculated based on Simpson’s (1949) in Excel.

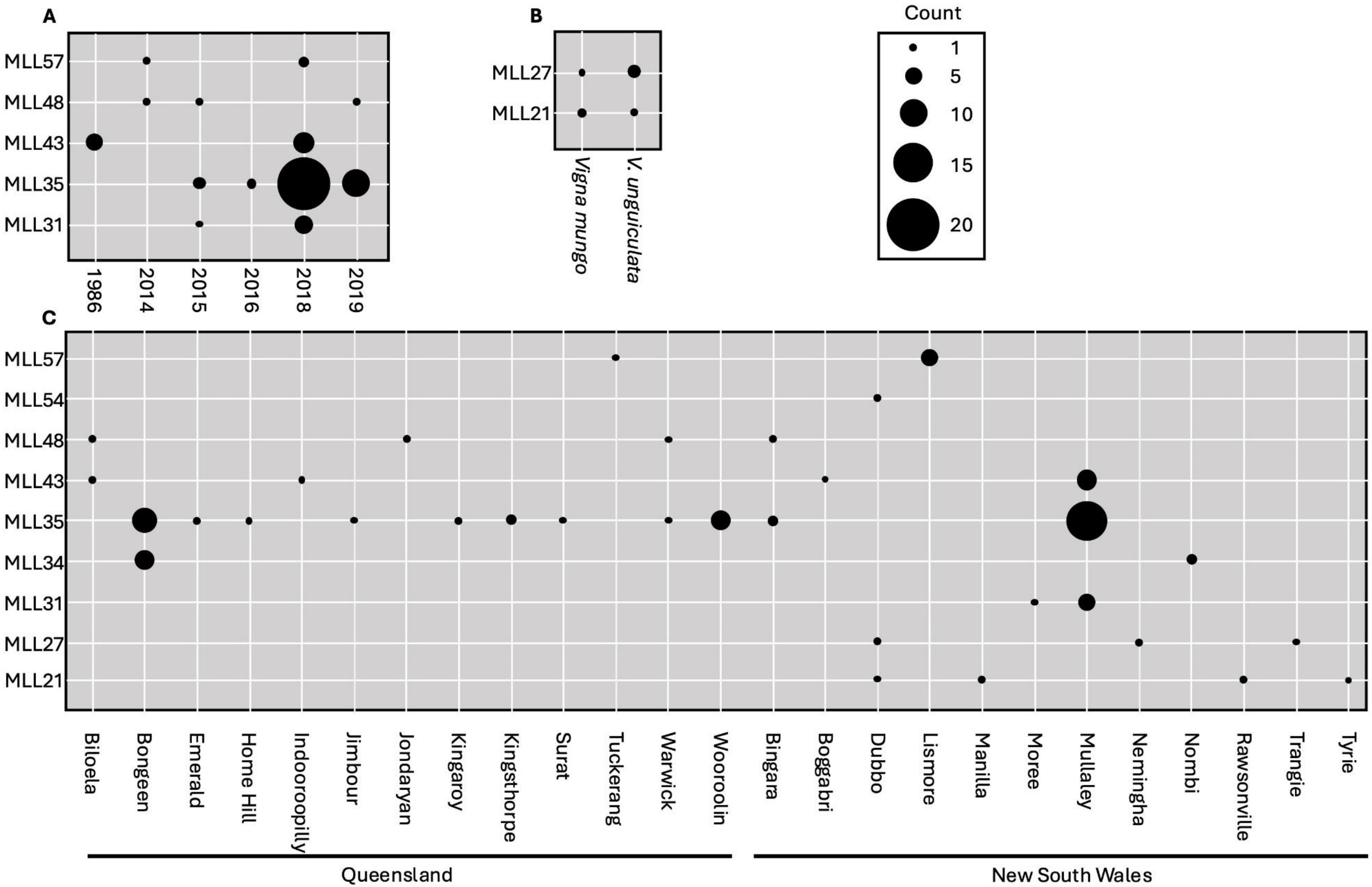
A
B
Count
1
5
10
15
20
MLL57
MLL48
MLL43
MLL35
MLL31
1986
2014
2015
2016
2018
2019
MLL27
MLL21
Vigna mungo
V. unguiculata
C
MLL57
MLL54
MLL48
MLL43
MLL35
MLL34
MLL31
MLL27
MLL21
Biloela
Bongeen
Emerald
Home Hill
Indooroopilly
Jimbour
Jondaryan
Kingaroy
Kingsthorpe
Surat
Tuckerang
Warwick
Wooroolin
Bingara
Boggabri
Dubbo
Lismore
Manilla
Moree
Mullaley
Nemingha
Nombi
Rawsonville
Trangie
Tyrie
Queensland
New South Wales

41

**Fig. S3.** Recurrent multi-locus lineages (MLLs) of *Curtobacterium flaccumfaciens* pv. *flaccumfaciens* shared among sampled (A) years, (B) hosts, and (C) locations. Circles represent MLLs shown on the vertical axis, with circle sizes proportional to MLL frequencies depicted in the legend.

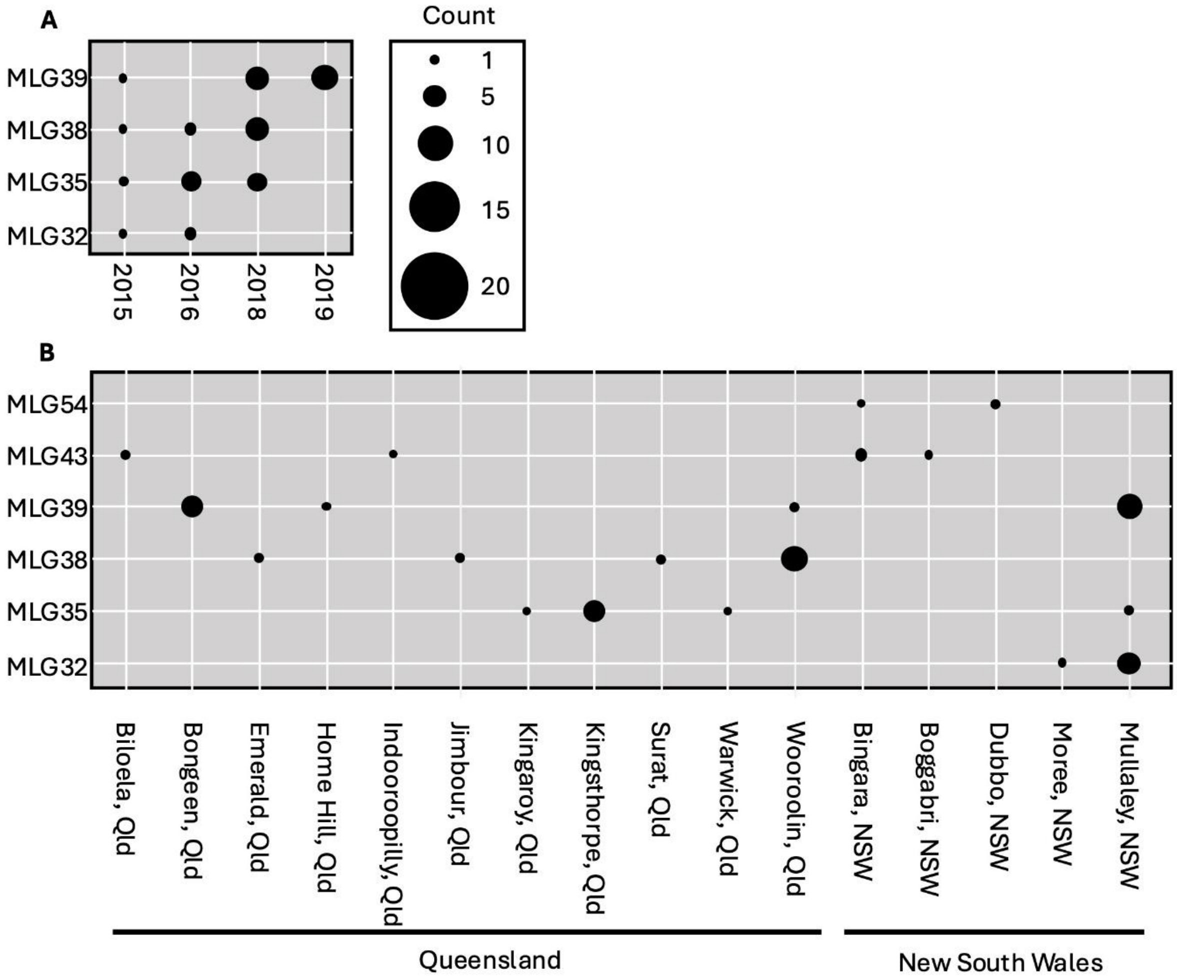


**Fig. S4.** Recurrent multi-locus genotypes (MLGs) of *Curtobacterium flaccumfaciens* pv. *flaccumfaciens* shared among sampled (A) years and (B) locations. Circles represent MLGs as shown on the vertical axis, with circle sizes proportional to MLG frequencies as depicted in the legend.

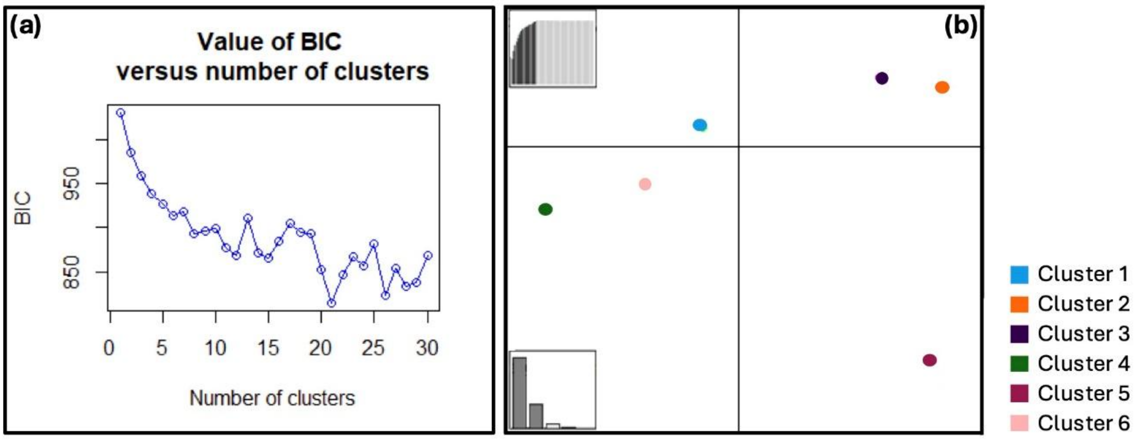


**Fig. S5.** Discriminant analysis of principal components (DAPC) for *Curtobacterium flaccumfaciens* pv. *flaccumfaciens* collected from 1986 to 2019 from various host crops in Australia. The graph on the left shows Bayesian information criterion (BIC) plotted against the number of inferred clusters for clone-corrected data set showed an increase in BIC after K = 6, which was selected as the number of clusters. The ordination plot on the right shows the six assigned genetically related clusters of individuals, which correspond to the clusters identified in the STRUCTURE analysis (Pritchard et al. 2000).

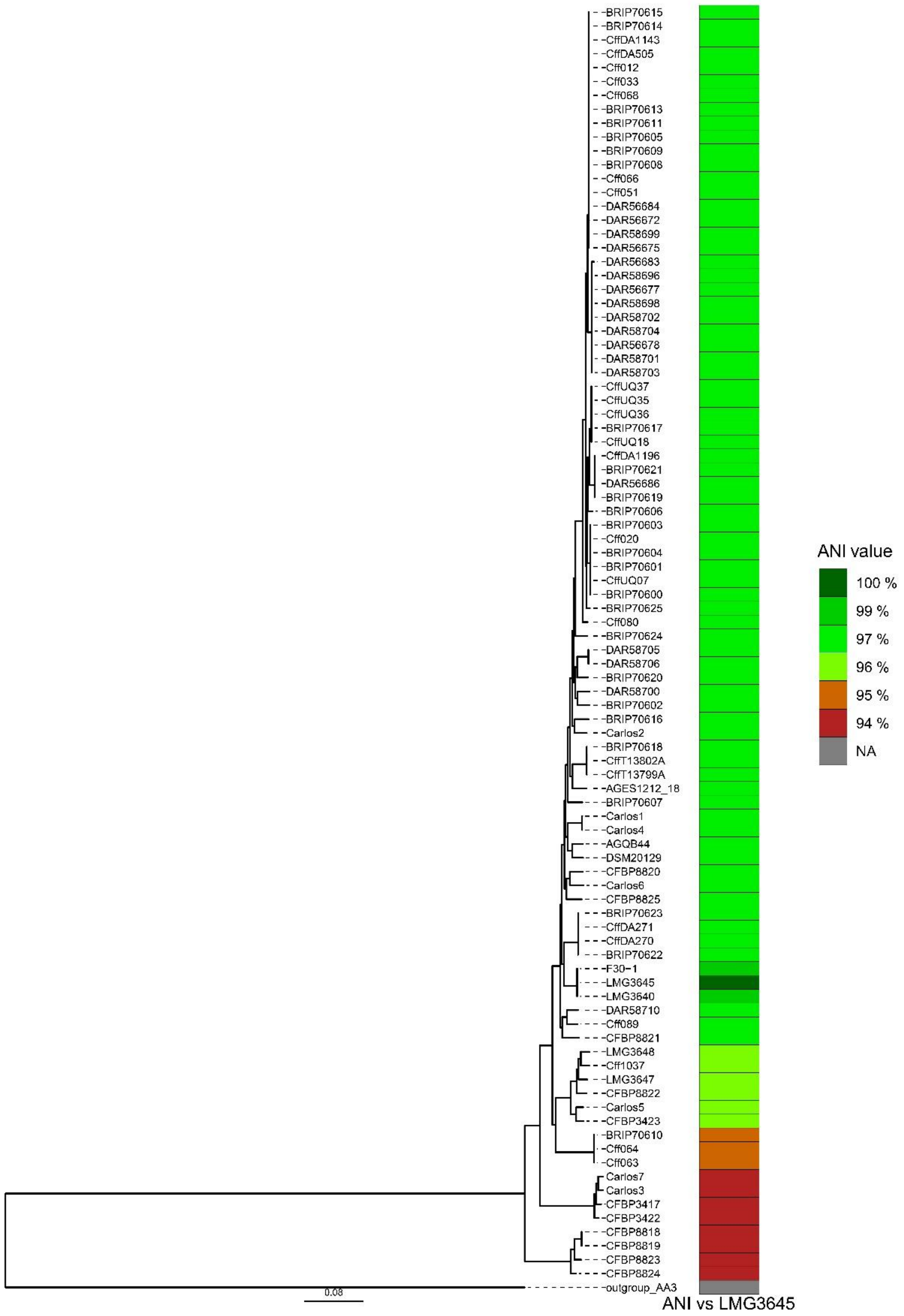


**Fig S6.** Maximum likelihood phylogeny of *Curtobacterium flaccumfaciens* pv. *flaccumfaciens* unique MLGs after clone correction produced in RAxML (Stamatakis 2014) based on 1,669 single-copy core genes. *Curtobacterium pusillum* AA3 (CP018783.1, CP018784.1) was used to root the tree. ANI scores were calculated using the *C. flaccumfaciens* pv. *flaccumfaciens* type strain LMG 3645 as referenced with FastANI (Jain et al. 2018).

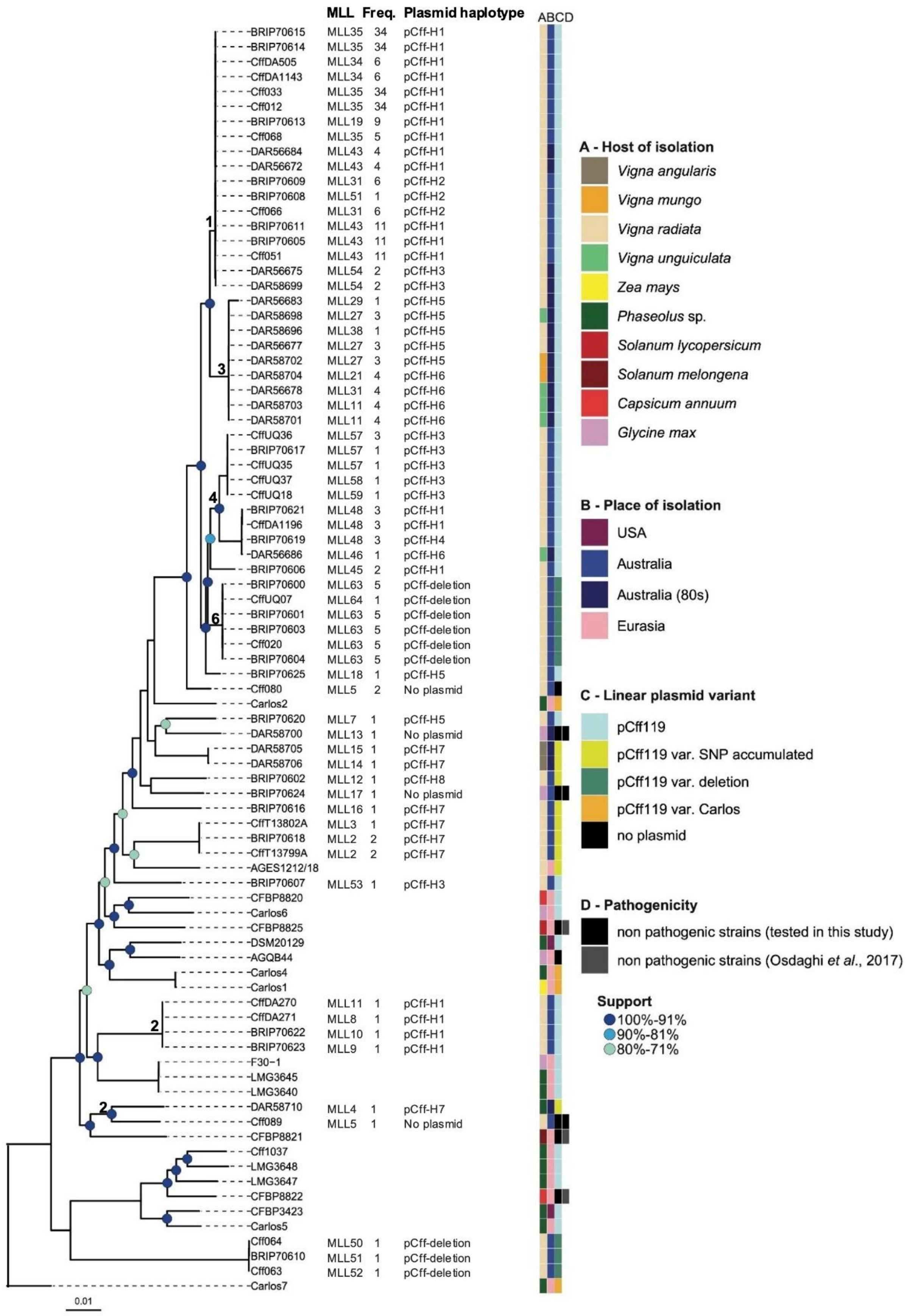
MLL Freq. Plasmid haplotype
ABCD
BRIP70615 MLL35 34 pCff-H1
BRIP70614 MLL35 34 pCff-H1
CffDA505 MLL34 6 pCff-H1
CffDA1143 MLL34 6 pCff-H1
Cff033 MLL35 34 pCff-H1
Cff012 MLL35 34 pCff-H1
BRIP70613 MLL19 9 pCff-H1
Cff068 MLL35 5 pCff-H1
DAR56684 MLL43 4 pCff-H1
DAR56672 MLL43 4 pCff-H1
BRIP70609 MLL31 6 pCff-H2
BRIP70608 MLL51 1 pCff-H2
Cff066 MLL31 6 pCff-H2
BRIP70611 MLL43 11 pCff-H1
BRIP70605 MLL43 11 pCff-H1
Cff051 MLL43 11 pCff-H1
DAR56675 MLL54 2 pCff-H3
DAR58699 MLL54 2 pCff-H3
DAR56683 MLL29 1 pCff-H5
DAR58698 MLL27 3 pCff-H5
DAR58696 MLL38 1 pCff-H5
DAR56677 MLL27 3 pCff-H5
DAR58702 MLL27 3 pCff-H5
DAR58704 MLL21 4 pCff-H6
DAR56678 MLL31 4 pCff-H6
DAR58703 MLL11 4 pCff-H6
DAR58701 MLL11 4 pCff-H6
CffUQ36 MLL57 3 pCff-H3
BRIP70617 MLL57 1 pCff-H3
CffUQ35 MLL57 1 pCff-H3
CffUQ37 MLL58 1 pCff-H3
CffUQ18 MLL59 1 pCff-H3
BRIP70621 MLL48 3 pCff-H1
CffDA1196 MLL48 3 pCff-H1
BRIP70619 MLL48 3 pCff-H4
DAR56686 MLL46 1 pCff-H6
BRIP70606 MLL45 2 pCff-H1
BRIP70600 MLL63 5 pCff-deletion
CffUQ07 MLL64 1 pCff-deletion
BRIP70601 MLL63 5 pCff-deletion
BRIP70603 MLL63 5 pCff-deletion
Cff020 MLL63 5 pCff-deletion
BRIP70604 MLL63 5 pCff-deletion
BRIP70625 MLL18 1 pCff-H5
Cff080 MLL5 2 No plasmid
Carlos2
BRIP70620 MLL7 1 pCff-H5
DAR58700 MLL13 1 No plasmid
DAR58705 MLL15 1 pCff-H7
DAR58706 MLL14 1 pCff-H7
BRIP70602 MLL12 1 pCff-H8
BRIP70624 MLL17 1 No plasmid
BRIP70616 MLL16 1 pCff-H7
CffT13802A MLL3 1 pCff-H7
BRIP70618 MLL2 2 pCff-H7
CffT13799A MLL2 2 pCff-H7
AGES1212/18
BRIP70607 MLL53 1 pCff-H3
CFBP8820
Carlos6
CFBP8825
DSM20129
AGQB44
Carlos4
Carlos1
CffDA270 MLL11 1 pCff-H1
CffDA271 MLL8 1 pCff-H1
BRIP70622 MLL10 1 pCff-H1
BRIP70623 MLL9 1 pCff-H1
F30-1
LMG3645
LMG3640
DAR58710 MLL4 1 pCff-H7
Cff089 MLL5 1 No plasmid
CFBP8821
Cff1037
LMG3648
LMG3647
CFBP8822
CFBP3423
Carlos5
Cff064 MLL50 1 pCff-deletion
BRIP70610 MLL51 1 pCff-deletion
Cff063 MLL52 1 pCff-deletion
Carlos7
1
2
3
4
6
2
0.01
A - Host of isolation
Vigna angularis
Vigna mungo
Vigna radiata
Vigna unguiculata
Zea mays
Phaseolus sp.
Solanum lycopersicum
Solanum melongena
Capsicum annuum
Glycine max
B - Place of isolation
USA
Australia
Australia (80s)
Eurasia
C - Linear plasmid variant
pCff119
pCff119 var. SNP accumulated
pCff119 var. deletion
pCff119 var. Carlos
no plasmid
D - Pathogenicity
non pathogenic strains (tested in this study)
non pathogenic strains (Osdaghi et al., 2017)
Support
100%-91%
90%-81%
80%-71%

**Fig. S7.** Maximum likelihood phylogeny of a global population of *Curtobacterium flaccumfaciens* pv. *flaccumfaciens* produced in RAxML version 7.2.8 (Stamatakis 2014) based on nucleotide sequences of 1,835 core genes after removal of recombinant regions. Carlos7 was used as outgroup. Clades numbers correspond to the Clusters detected by the Structure software. Isolates from Clusters 1, 3, 4, and 6 formed monophyletic groups while isolates from Clusters 2 and 5 were interspersed among *C. flaccumfaciens* pv. *flaccumfaciens* isolates from different hosts and countries. MLLs represent unique multi-locus lineages (MLLs) (Supplementary Table S1). While four general categories of plasmid variants are defined for the entire dataset (pCff119, pCff199 SNP accumulated, pCff119 deletion, pCff119 Carlos), plasmid haplotypes of the Australian isolates (based on 99 plasmid SNPs) from the haplotypic network in Fig. 4 are also provided next to the Australian isolates. Scale bar indicates nucleotide substitutions per site.

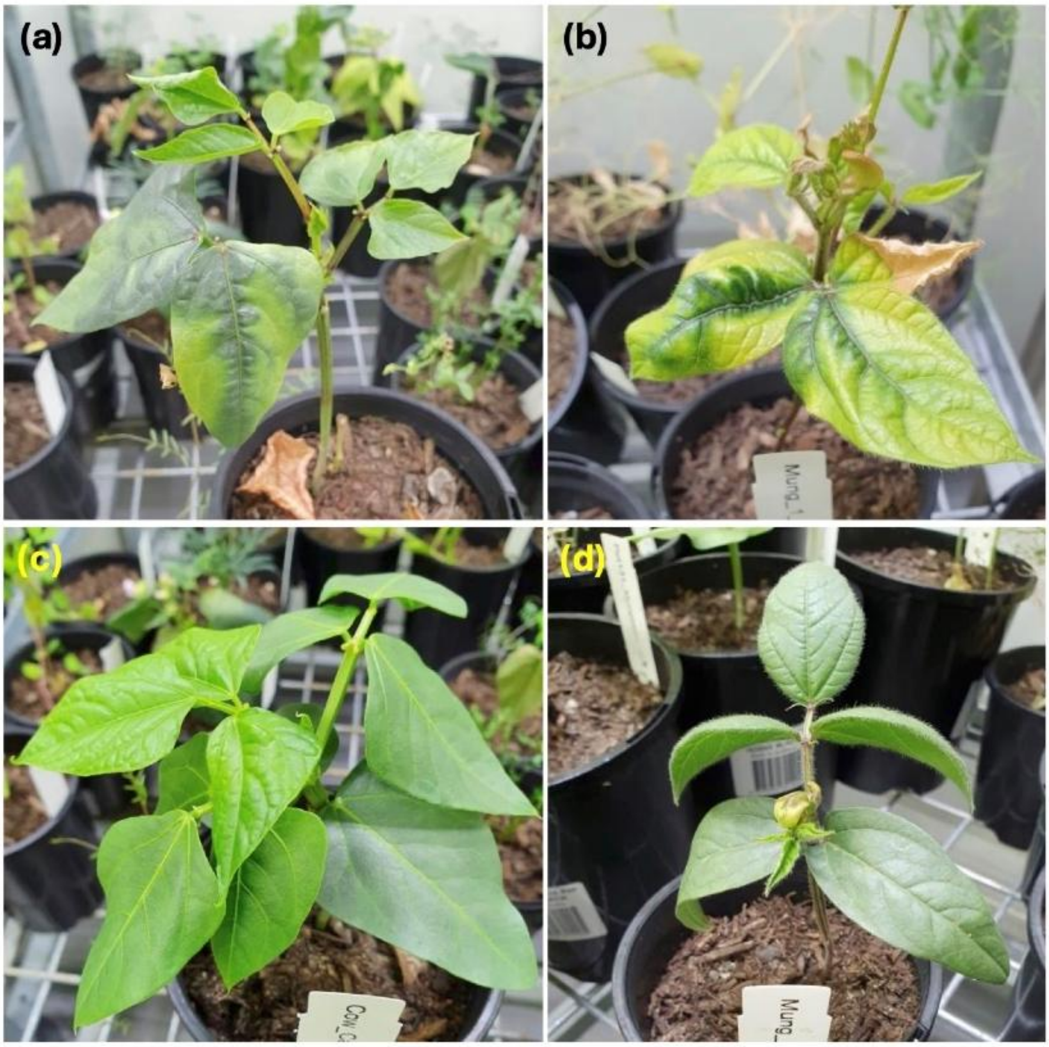


**Fig. S8.** Tan spot symptoms caused by *Curtobacterium flaccumfaciens* pv. *flaccumfaciens* isolate BRIP 70623 on **(a)** cowpea (cv. Red Caloona) and **(b)** mungbean (cv. Jade-AU) at 25 days after inoculation. Inoculation of **(c)** cowpea (cv. Red Caloona) and **(d)** mungbean (cv. Jade-AU) by isolate BRIP 70624, which lack a linear plasmid, did not result in any symptoms.

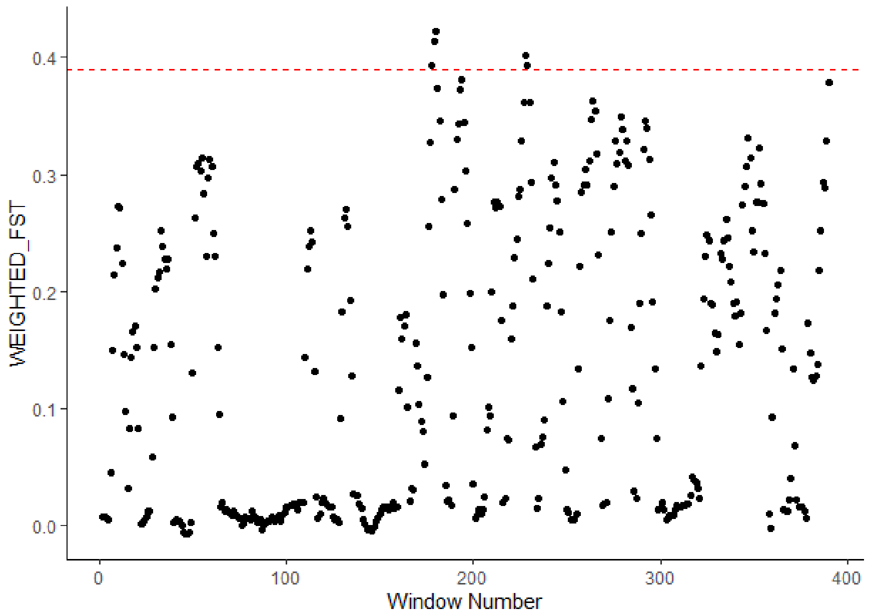


**Fig S9.** Manhattan plot illustrating SNP-by-SNP $F_{st}$ values across the genome, comparing historical (1986-87) and contemporary (2014-2019) populations of *Curtobacterium flaccumfaciens* pv. *flaccumfaciens*. $F_{st}$ values were calculated using a sliding window approach (50,000 bp window size, 50,000 bp step) to identify loci potentially under selection. The horizontal dashed line represents the 99th percentile threshold, with SNPs on the windows above this line classified as outliers, indicating potential loci under selection.

**Table S1.** *Curtobacterium flaccumfaciens* pv. *flaccumfaciens* isolates (*n* = 119) collected from mungbean growing regions in Australia. Only 117 isolates were included in the population genomic analyses using whole genome sequence data as the isolates depicted in bold were removed from the analyses due to > 20% missing data. MLGs and MLLs were identified based on 43,974 genome wide single nucleotide polymorphisms. Clusters are assigned based on the structure analysis using the software STRUCTURE (Pritchard et al. 2000), where admixed isolates represent those that could not be assigned to a cluster based on probability of > 0.6.

| **Isolate** | **MLG** | **MLL** | **Cluster** | **Year** | **State** | **Field** | **Plant** | **Leaf** | **Location** | **Host** |
|---|---|---|---|---|---|---|---|---|---|---|
| Cff002 | MLG61 | MLL63 | 6 | 2018 | NSW | 42 | P1 | L2 | Mullaley | *Vigna radiata* |
| Cff003; BRIP 70600 | MLG66 | MLL63 | 6 | 2018 | NSW | 42 | P1 | L3 | Mullaley | *V. radiata* |
| Cff007 | MLG32 | MLL31 | 1 | 2018 | NSW | 42 | P1 | L6 | Mullaley | *V. radiata* |
| Cff009 | MLG39 | MLL35 | 1 | 2018 | NSW | 42 | P5 | L10 | Mullaley | *V. radiata* |
| Cff010 | MLG39 | MLL35 | 1 | 2018 | NSW | 42 | P6 | L11 | Mullaley | *V. radiata* |
| Cff012 | MLG37 | MLL35 | 1 | 2018 | NSW | 42 | P7 | L12 | Mullaley | *V. radiata* |
| Cff014; BRIP 70601 | MLG65 | MLL63 | 6 | 2018 | NSW | 42 | P9 | L14 | Mullaley | *V. radiata* |
| Cff016; BRIP 70602 | MLG12 | MLL12 | Admixed (2&5) | 2018 | NSW | 42 | P11 | L16 | Mullaley | *V. radiata* |
| Cff018 | MLG42 | MLL43 | 1 | 2018 | NSW | 42 | P12 | L17 | Mullaley | *V. radiata* |
| Cff019; BRIP 70603 | MLG63 | MLL63 | 6 | 2018 | NSW | 42 | P13 | L18 | Mullaley | *V. radiata* |
| Cff020 | MLG60 | MLL63 | 6 | 2018 | NSW | 42 | P13 | L18 | Mullaley | *V. radiata* |
| Cff021; BRIP 70604 | MLG61 | MLL63 | 6 | 2018 | NSW | 42 | P14 | L19 | Mullaley | *V. radiata* |
| Cff026 | MLG37 | MLL35 | 1 | 2018 | NSW | 42 | P21 | L21 | Mullaley | *V. radiata* |
| Cff029; BRIP 70605 | MLG42 | MLL43 | 1 | 2018 | NSW | 42 | P21 | L22 | Mullaley | *V. radiata* |
| Cff033 | MLG36 | MLL35 | 1 | 2018 | NSW | 42 | P21 | L24 | Mullaley | *V. radiata* |
| Cff034 | MLG36 | MLL35 | 1 | 2018 | NSW | 42 | P21 | L24 | Mullaley | *V. radiata* |
| Cff035 | MLG36 | MLL35 | 1 | 2018 | NSW | 42 | P21 | L24 | Mullaley | *V. radiata* |
| Cff037; BRIP 70606 | MLG45 | MLL45 | Admixed (4&6) | 2018 | NSW | 42 | P21 | L25 | Mullaley | *V. radiata* |

| Isolate | MLG | MLL | Cluster | Year | State | Field | Plant | Leaf | Location | Host |
|---|---|---|---|---|---|---|---|---|---|---|
| Cff040 | MLG44 | MLL45 | Admixed (4&6) | 2018 | NSW | 42 | P21 | L25 | Mullaley | *V. radiata* |
| Cff043; BRIP 70607 | MLG53 | ML53 | Admixed (2&5) | 2018 | NSW | 42 | P21 | L26 | Mullaley | *V. radiata* |
| Cff051 | MLG41 | MLL43 | 1 | 2018 | NSW | 42 | P25 | L30 | Mullaley | *V. radiata* |
| Cff052; BRIP 70608 | MLG32 | MLL31 | 1 | 2018 | NSW | 42 | P26 | L31 | Mullaley | *V. radiata* |
| Cff054 | MLG36 | MLL35 | 1 | 2018 | NSW | 42 | P27 | L32 | Mullaley | *V. radiata* |
| Cff055 | MLG36 | MLL35 | 1 | 2018 | NSW | 42 | P27 | L32 | Mullaley | *V. radiata* |
| Cff056 | MLG36 | MLL35 | 1 | 2018 | NSW | 42 | P27 | L32 | Mullaley | *V. radiata* |
| Cff058 | MLG37 | MLL35 | 1 | 2018 | NSW | 42 | P28 | L33 | Mullaley | *V. radiata* |
| Cff059 | MLG31 | MLL31 | 1 | 2018 | NSW | 42 | P29 | L34 | Mullaley | *V. radiata* |
| Cff061; BRIP 70609 | MLG31 | MLL31 | 1 | 2018 | NSW | 42 | P29 | L34 | Mullaley | *V. radiata* |
| Cff062; BRIP 70610 | MLG51 | MLL51 | 2 | 2018 | NSW | 42 | P30 | L35 | Mullaley | *V. radiata* |
| Cff063 | MLG52 | MLL52 | 2 | 2018 | NSW | 42 | P30 | L35 | Mullaley | *V. radiata* |
| Cff064 | MLG50 | MLL50 | 2 | 2018 | NSW | 42 | P30 | L35 | Mullaley | *V. radiata* |
| Cff066 | MLG30 | MLL31 | 1 | 2018 | NSW | 42 | P32 | L37 | Mullaley | *V. radiata* |
| Cff068 | MLG35 | MLL35 | 1 | 2018 | NSW | 42 | P33 | L38 | Mullaley | *V. radiata* |
| Cff069 | MLG42 | MLL43 | 1 | 2018 | NSW | 42 | P34 | L39 | Mullaley | *V. radiata* |
| Cff070 | MLG42 | MLL43 | 1 | 2018 | NSW | 42 | P34 | L39 | Mullaley | *V. radiata* |
| Cff071; BRIP 70611 | MLG42 | MLL43 | 1 | 2018 | NSW | 42 | P34 | L39 | Mullaley | *V. radiata* |
| Cff072 | MLG39 | MLL35 | 1 | 2018 | NSW | 42 | P35 | L40 | Mullaley | *V. radiata* |
| Cff080 | MLG6 | MLL5 | Admixed (6&4&5) | 2018 | Qld | 10 | P8 | L13 | Brookstead | *V. radiata* |
| Cff089 | MLG5 | MLL5 | Admixed (2&5) | 2018 | Qld | 10 | P4 | L29 | Brookstead | *V. radiata* |
| Cff101 | MLG19 | MLL19 | 1 | 2018 | Qld | 10 | P14 | L39 | Brookstead | *V. radiata* |
| Cff102 | MLG19 | MLL19 | 1 | 2018 | Qld | 10 | P14 | L39 | Brookstead | *V. radiata* |
| Cff103 | MLG20 | MLL19 | 1 | 2018 | Qld | 10 | P14 | L39 | Brookstead | *V. radiata* |

| Isolate | MLG | MLL | Cluster | Year | State | Field | Plant | Leaf | Location | Host |
|---|---|---|---|---|---|---|---|---|---|---|
| Cff106 | MLG19 | MLL19 | 1 | 2018 | Qld | 10 | P10 | L35 | Brookstead | *V. radiata* |
| Cff107; BRIP 70612 | MLG19 | MLL19 | 1 | 2018 | Qld | 10 | P10 | L35 | Brookstead | *V. radiata* |
| Cff108 | MLG19 | MLL19 | 1 | 2018 | Qld | 10 | P11 | L36 | Brookstead | *V. radiata* |
| Cff109 | MLG19 | MLL19 | 1 | 2018 | Qld | 10 | P11 | L36 | Brookstead | *V. radiata* |
| Cff110; BRIP 70613 | MLG19 | MLL19 | 1 | 2018 | Qld | 10 | P12 | L37 | Brookstead | *V. radiata* |
| Cff111 | MLG19 | MLL19 | 1 | 2018 | Qld | 10 | P12 | L37 | Brookstead | *V. radiata* |
| Cff118 | MLG39 | MLL35 | 1 | 2018 | Qld | 58 | - | - | Wooroolin | *V. radiata* |
| Cff129 | MLG38 | MLL35 | 1 | 2018 | Qld | 58 | - | - | Wooroolin | *V. radiata* |
| Cff130; BRIP 70614 | MLG38 | MLL35 | 1 | 2018 | Qld | 58 | - | - | Wooroolin | *V. radiata* |
| Cff192 | MLG38 | MLL35 | 1 | 2018 | Qld | 58 | - | - | Wooroolin | *V. radiata* |
| Cff199 | MLG38 | MLL35 | 1 | 2018 | Qld | 58 | - | - | Wooroolin | *V. radiata* |
| Cff200; BRIP 70615 | MLG38 | MLL35 | 1 | 2018 | Qld | 58 | - | - | Wooroolin | *V. radiata* |
| **CffDA027** | - | - | - | - | Qld | 1436 | - | - | Missen Flat | *V. radiata* |
| CffDA1121; BRIP 70625 | MLG18 | MLL18 | Admixed (1&4&6) | 2019 | Qld | 10 | P1 | L1 | Formartin | *V. radiata* |
| CffDA1143 | MLG34 | MLL34 | 1 | 2019 | Qld | 10 | P1 | L2 | Bongeen | *V. radiata* |
| CffDA1145 | MLG34 | MLL34 | 1 | 2019 | Qld | 10 | P1 | L3 | Bongeen | *V. radiata* |
| CffDA1149 | MLG34 | MLL34 | 1 | 2019 | Qld | 10 | P2 | L1 | Bongeen | *V. radiata* |
| CffDA1150 | MLG34 | MLL34 | 1 | 2019 | Qld | 23 | - | - | Bongeen | *V. radiata* |
| CffDA1152 | MLG39 | MLL35 | 1 | 2019 | Qld | 45 | - | - | Bongeen | *V. radiata* |
| CffDA1153 | MLG39 | MLL35 | 1 | 2019 | Qld | 45 | - | - | Bongeen | *V. radiata* |
| CffDA1155 | MLG39 | MLL35 | 1 | 2019 | Qld | 52 | - | - | Bongeen | *V. radiata* |
| CffDA1156 | MLG39 | MLL35 | 1 | 2019 | Qld | 52 | - | | Bongeen | *V. radiata* |
| CffDA1166 | MLG35 | MLL35 | 1 | 2019 | Qld | 40 | - | - | Kingsthorpe | *V. radiata* |
| CffDA1169 | MLG35 | MLL35 | 1 | 2019 | Qld | 255 | - | - | Kingsthorpe | *V. radiata* |

| Isolate | MLG | MLL | Cluster | Year | State | Field | Plant | Leaf | Location | Host |
|---|---|---|---|---|---|---|---|---|---|---|
| CffDA1196 | MLG48 | MLL48 | 4 | 2019 | Qld | 267 | - | - | Jondaryan | *V. radiata* |
| CffDA270 | MLG11 | MLL11 | 2 | 2019 | Qld | 267 | - | - | Brookstead | *V. radiata* |
| CffDA271 | MLG8 | MLL8 | 2 | 2019 | Qld | 267 | - | - | Brookstead | *V. radiata* |
| CffDA272; BRIP 70622 | MLG10 | MLL10 | 2 | 2019 | Qld | 267 | - | - | Brookstead | *V. radiata* |
| CffDA274; BRIP 70623 | MLG9 | MLL9 | 2 | 2019 | Qld | 267 | - | - | Brookstead | *V. radiata* |
| CffDA492 | MLG39 | MLL35 | 1 | 2019 | Qld | 267 | - | - | Bongeen | *V. radiata* |
| CffDA493 | MLG39 | MLL35 | 1 | 2019 | NSW | 267 | - | - | Mullaley | *V. radiata* |
| CffDA494 | MLG39 | MLL35 | 1 | 2019 | NSW | 267 | - | - | Mullaley | *V. radiata* |
| CffDA505 | MLG33 | MLL34 | 1 | 2019 | NSW | 258 | - | - | Nombi | *V. radiata* |
| CffDA506 | MLG33 | MLL34 | 1 | 2019 | NSW | 258 | - | - | Nombi | *V. radiata* |
| CffDA535; BRIP 70624 | MLG17 | MLL17 | 5 | 2019 | NSW | 295 | - | - | Sherwood | *Glycine max* |
| CffT13692; BRIP 70616 | MLG16 | MLL16 | 5 | 2014 | Qld | T13692 | - | - | Tuckerang | *V. radiata* |
| CffT13694; BRIP 70617 | MLG56 | MLL57 | 4 | 2014 | Qld | T13694 | - | - | Tuckerang | *V. radiata* |
| CffT13799A | MLG2 | MLL2 | 5 | 2014 | Qld | T13799 | - | - | Warwick | *V. radiata* |
| CffT13799B; BRIP 70618 | MLG1 | MLL2 | 5 | 2014 | Qld | T13799 | - | - | Warwick | *V. radiata* |
| CffT13802A | MLG3 | MLL3 | 5 | 2014 | Qld | T13802 | - | - | Warwick | *V. radiata* |
| CffT13802B; BRIP 70619 | MLG49 | MLL48 | 4 | 2014 | Qld | T13802 | - | - | Warwick | *V. radiata* |
| CffT13932 | MLG32 | MLL31 | 1 | 2015 | NSW | T13932 | - | - | Moree | *V. radiata* |
| CffT14011 | MLG38 | MLL35 | 1 | 2015 | Qld | T14069 | - | - | Surat | *V. radiata* |
| CffT14069; BRIP 70620 | MLG7 | MLL7 | 5 | 2015 | Qld | T14069 | - | - | Ayr | *V. radiata* |
| CffT14074 | MLG38 | MLL35 | 1 | 2015 | Qld | T14074 | - | - | Emerald | *V. radiata* |
| CffT14089; BRIP 70621 | MLG47 | MLL48 | 4 | 2015 | Qld | T14089 | - | - | Biloela | *V. radiata* |
| CffT14103 | MLG39 | MLL35 | 1 | 2015 | Qld | T14103 | - | - | Home Hill | *V. radiata* |
| CffT14141 | MLG39 | MLL35 | 1 | 2016 | Qld | T14141 | - | - | Jimbour | *V. radiata* |

| Isolate | MLG | MLL | Cluster | Year | State | Field | Plant | Leaf | Location | Host |
|---|---|---|---|---|---|---|---|---|---|---|
| CffUQ07 | MLG64 | MLL64 | 6 | 2016 | Qld | Unknown1 | - | - | Canaga | *V. radiata* |
| CffUQ18 | MLG59 | MLL59 | 4 | 2016 | Qld | Unknown2 | - | - | Kingaroy | *V. radiata* |
| **CffUQ19** | - | - | - | 2016 | Qld | Unknown2 | - | - | Kingaroy | *V. radiata* |
| CffUQ20 | MLG35 | MLL35 | 1 | 2016 | Qld | Unknown2 | - | - | Kingaroy | *V. radiata* |
| CffUQ26 | MLG35 | MLL35 | 1 | 2018 | Qld | Unknown3 | - | - | Warwick | *V. radiata* |
| CffUQ35 | MLG55 | MLL57 | 4 | 2018 | NSW | Unknown4 | - | - | Lismore | *V. radiata* |
| CffUQ36 | MLG57 | MLL57 | 4 | 2018 | NSW | Unknown4 | - | - | Lismore | *V. radiata* |
| CffUQ37 | MLG58 | MLL58 | 4 | 2018 | NSW | Unknown4 | - | - | Lismore | *V. radiata* |
| DAR 56672 | MLG43 | MLL43 | 1 | 1986 | Qld | Unknown | - | - | Biloela | *V. radiata* |
| DAR 56673 | MLG43 | MLL43 | 1 | 1986 | Qld | Unknown | - | - | Indooroopilly | *V. radiata* |
| DAR 56675 | MLG54 | MLL54 | 1 | 1987 | NSW | Unknown | - | - | Bingara | *V. radiata* |
| DAR 56677 | MLG26 | MLL27 | 3 | 1987 | NSW | Unknown | - | - | Nemingha | *V. mungo* |
| DAR 56678 | MLG24 | MLL21 | 3 | 1987 | NSW | Unknown | - | - | Manilla | *Vigna unguiculata* |
| DAR 56683 | MLG29 | MLL29 | Admixed (1&3) | 1986 | NSW | Unknown | - | - | Moree | *V. radiata* |
| DAR 56684 | MLG43 | MLL43 | 1 | 1986 | NSW | Unknown | - | - | Boggabri | *V. radiata* |
| DAR 56685 | MLG43 | MLL43 | 1 | 1986 | NSW | Unknown | - | - | Bingara | *V. radiata* |
| DAR 56686 | MLG46 | MLL46 | Admixed (1&4) | 1986 | NSW | Unknown | - | - | Tamworth | *V. unguiculata* |
| DAR 58696 | MLG28 | MLL38 | Admixed (1&3) | 1986 | NSW | Unknown | - | - | Bingara | *V. radiata* |
| DAR 58697 | MLG40 | MLL43 | 1 | 1986 | NSW | Unknown | - | - | Bingara | *V. radiata* |
| DAR 58698 | MLG27 | MLL27 | 3 | 1987 | NSW | Unknown | - | - | Dubbo | *V. unguiculata* |
| DAR 58699 | MLG54 | MLL54 | 1 | 1987 | NSW | Unknown | - | - | Dubbo | *V. radiata* |
| DAR 58700 | MLG13 | MLL13 | 5 | 1987 | NSW | Unknown | - | - | Dubbo | *G. max* |
| DAR 58701 | MLG21 | MLL21 | 3 | 1987 | NSW | Unknown | - | - | Dubbo | *V. unguiculata* |
| DAR 58702 | MLG25 | MLL27 | 3 | 1987 | NSW | Unknown | - | - | Trangie | *Vigna mungo* |

| Isolate | MLG | MLL | Cluster | Year | State | Field | Plant | Leaf | Location | Host |
|---|---|---|---|---|---|---|---|---|---|---|
| DAR 58703 | MLG23 | MLL21 | 3 | 1987 | NSW | Unknown | - | - | Rawsonville | *V. unguiculata* |
| DAR 58704 | MLG22 | MLL21 | 3 | 1987 | NSW | Unknown | - | - | Tyrie | *V. mungo* |
| DAR 58705 | MLG15 | MLL15 | 5 | 1987 | NSW | Unknown | - | - | Rawsonville | *Vigna angularis* |
| DAR 58706 | MLG14 | MLL14 | 5 | 1987 | NSW | Unknown | - | - | Trangie | *V. angularis* |
| DAR 58710 | MLG4 | MLL4 | 2 | 1987 | NSW | Unknown | - | - | Tamworth | *Phaseolus vulgaris* |

**Table S2.** Indices of genetic diversity for the Australian populations of *Curtobacterium flaccumfaciens* pv. *flaccumfaciens* collected from New South Wales (NSW) and Queensland (Qld), Australia, between 1986 and 2019.

| Population | N[a] | MLL[b] | eMLL[c] | SE[d] | λ[e] | CF[f] | $H_{exp}$[g] |
|---|---|---|---|---|---|---|---|
| NSW | 65 | 25 | 21.8 | 1.38 | 0.90 | 62% | 0.007 |
| Qld | 52 | 19 | 19.0 | 0.00 | 0.82 | 63% | 0.007 |
| Total | 117 | 40 | 23.4 | 2.32 | 0.89 | 66% | 0.007 |

[a] N = population size.

[b] MLL = Number of multi-locus lineages after contracting MLGs with one to seven SNP differences into one MLL.

[c] eMLL = expected number of MLLs at the lowest common sample size to allow comparison between populations (after rarefaction).

[d] SE = The standard error for the rarefaction analysis for eMLLs.

[e] λ = defined here as unbiased Simpson's (Simpson 1949) complement index of genotypic diversity, defined as the probability that two genotypes randomly chosen from the population are different, corrected for sample size.

[f] CF = Clonal fraction; N-number of MLLs)/N, where N is total number of isolates.

[g] $H_{exp}$ = Nei's gene diversity (expected heterozygosity) (Nei 1978).

**Table S3.** Indices of genetic diversity of the Australian *Curtobacterium flaccumfaciens* pv. *flaccumfaciens* isolates intensively sampled from field 42 in New South Wales and field 10 in Queensland, in 2019.

| **Field** | **N**[a] | **MLL**[b] | **eMLL**[c] | **SE**[d] | **λ**[e] | **E5**[f] | **CF**[g] | **Hexp**[g] |
|---|---|---|---|---|---|---|---|---|
| 42 | 37 | 11 | 6.9 | 1.17 | 0.81 | 0.70 | 70% | 0.006 |
| 10 | 15 | 7 | 7.0 | 0.00 | 0.71 | 0.53 | 53% | 0.003 |

[a] N = population size.

[b] MLL = Number of multi-locus lineages after contracting MLGs with one to seven SNP differences into one MLL.

[c] eMLL = expected number of MLLs at the lowest common sample size to allow comparison between populations (after rarefaction).

[d] SE = The standard error for the rarefaction analysis for eMLLs.

[e] λ = defined here as unbiased Simpson's (Simpson 1949) complement index of genotypic diversity, defined as the probability that two genotypes randomly chosen from the population are different, corrected for sample size.

[f] E5 = Evenness as a measure of the distribution of MLL abundances, where in a population with equally abundant MLLs, E5 = 1, and in a population dominated by a single MLL, E5=0.

[g] CF = Clonal fraction; N-number of MLLs)/N, where N is total number of isolates.

[h] $H_{exp}$ = Nei's gene diversity (expected heterozygosity) (Nei 1978).

**Table S4.** Private single nucleotide polymorphisms (SNPs) specific to *Curtobacterium flaccumfaciens* pv. *flaccumfaciens* MLL35, the most frequently sampled multi-locus lineage in Australian mungbean fields from 1986 to 2019.

| Position on the genome (bp) | REF[a] | ALT[b] | AF[c] | Annotation[d] |
|---|---|---|---|---|
| 69,834 | C | A | 0.05 | Stop gained, High impact, cation:dicarboxylase symporter family transporter; Loss of function (LOF) and nonsense-mediated decay (NMD) predicted. |
| 261,181 | T | A | 0.05 | Missense variant, Moderate impact, ROK (repressor, open reading frame, kinase) family transcriptional regulator |
| 704,799 | C | T | 0.05 | Missense variant, Moderate impacts, NAD-dependent epimerase/dehydratase family protein |
| 864,942 | C | G | 0.05 | Synonymous variant, Low impact, iolC (2-deoxy-5-keto-d-gluconate kinase) |
| 1,088,029 | A | G | 0.07 | Synonymous variant, Low impact, ABC transporter permease |
| 1,346,754 | C | G | 0.07 | Missense variant, Moderate impact, hypothetical protein |
| 1,673,380 | T | C | 0.05 | Missense variant, Moderate impact, aldo/keto reductase |
| 2,099,272 | G | A | 0.05 | Missense variant, Moderate impact, gltB Glutamate synthase |
| 3,511,809 | C | T | 0.05 | Missense variant, Moderate impact, hypothetical protein |
| 3,570,887 | T | C | 0.05 | Missense variant, Moderate impact, NCS2 family permease |

[a] REF = reference allele

[b] ALT = Alternate allele

[c] AF = Alelle frequency within the *Curtobacterium flaccumfaciens* pv. *flaccumfaciens* population

[d] Annotation of SNPs against gene models of the BRIP 70614 reference genome using SnpEff (Cingolani et al. 2012) which identifies variations with potential high, moderate, low, or modifier (intergenic) impact.

**Table S5.** Outlier loci detected through pairwise comparison of fixation index ($F_{st}$) between the historical and contemporary populations of *Curtobacterium flaccumfaciens* pv. *flaccumfaciens* in Australia.

| Position on the genome (bp) | GO[a] | Annotation[b] |
|---|---|---|
| 1,779,930 | glnA hypothetical type I glutamate-ammonia ligase | downstream gene variant |
| 1,780,188 | DUF4191 domain-containing protein | synonymous variant, low impact |
| 1,781,044 | Hypothetical protein | missense variant, moderate impact |
| 1,781,743 | Hypothetical protein | missense variant, moderate impact |
| 1,782,074 | GO:0016746 - acyltransferase activity (sucB hypothetical 2-oxoglutarate dehydrogenase encoding gene) | synonymous variant, low impact |
| 1,782,242 | GO:0016746 - acyltransferase activity (sucB hypothetical 2-oxoglutarate dehydrogenase encoding gene) | synonymous variant, low impact |
| 1,784,655 | GO:0004148 - dihydrolipoyl dehydrogenase activity (lpdA hypothetical dihydrolipoyl dehydrogenase encoding gene) | synonymous variant, low impact |
| 1,785,825 | GO:0030145 - manganese ion binding (leucyl aminopeptidase encoding gene) | synonymous variant, low impact |
| 1,785,840 | GO:0030145 - manganese ion binding (leucyl aminopeptidase encoding gene) | synonymous variant, low impact |
| 1,786,734 | PAC2 family protein, proteasome assembly chaperone family encoding gene | synonymous variant, low impact |
| 1,818,205 | Ribonuclease D | upstream gene variant |
| 2,294,743 | Hypothetical protein | missense variant, moderate impact |
| 2,296,400 | GO:0030170 - pyridoxal phosphate binding (cysteine desulfurase encoding gene) | synonymous variant, low impact |
| 2,299,068 | GO:0003824 - catalytic activity (glycogen debranching enzyme encoding gene) | synonymous variant, low impact |
| 2,308,041 | Hypothetical protein | missense variant, moderate impact |
| 2,309,119 | Transglycosylase SLT domain containing protein | synonymous variant, low impact |
| 2,310,105 | Transglycosylase SLT domain containing protein | synonymous variant, low impact |
| 2,310,301 | Transglycosylase SLT domain containing protein | upstream gene variant |
| 2,310,438 | Transglycosylase SLT domain containing protein | upstream gene variant |
| 2,311,236 | DivIVA domain containing protein | missense variant, moderate impact |
| 2,311,961 | GO:0004605 - phosphatidate cytidylyltransferase activity | synonymous variant, low impact |
| 2,312,090 | GO:0004605 - phosphatidate cytidylyltransferase activity | synonymous variant, low impact |
| 2,313,801 | GO:0033862 - UMP kinase activity | synonymous variant, low impact |
| 2,317,259 | GO:0022857 - transmembrane transporter activity (sugar porter family MFS transporter) | synonymous variant, low impact |
| 2,317,955 | GO:0008135 - translation factor activity, RNA binding | upstream gene variant |
| 2,318,079 | GO:0046872 - metal ion binding (M23_family_metallopeptidase - murein hydrolase activator EnvC) | missense variant, moderate impact |

| Position on the genome (bp) | GO[a] | Annotation[b] |
|---|---|---|
| 2,318,576 | GO:0046872 - metal ion binding (M23_family_metallopeptidase - murein hydrolase activator EnvC) | missense variant, moderate impact |
| 2,319,250 | GO:0003677 - DNA binding (tyrosine_recombinase XerC) | synonymous variant, low impact |
| 2,319,388 | GO:0003677 - DNA binding (tyrosine_recombinase XerC) | synonymous variant, low impact |
| 2,319,968 | GO:0009294 - DNA-mediated transformation (DNA-processing protein DprA) | synonymous variant, low impact |

[a] Gene Ontology

[b] Annotation of SNPs against gene models of the BRIP 70614 reference genome using SnpEff (Cingolani et al. 2012) which identifies variations with potential high, moderate, low, or modifier (intergenic) impact.

**Table S6.** *Curtobacterium flaccumfaciens* pv. *flaccumfaciens* reference genomes retrieved from National Centre for Biotechnology information (NCBI) database in December 2023.

| **Strain** | **Country of origin** | **Host of isolation** | **Year of isolation** | **NCBI accession** |
|---|---|---|---:|---|
| AGES1212/18 | Belgium | *Vigna radiata* | 2020 | JANVAN01 |
| AGQB44 | Belgium | *Glycine max* | 2022 | CP104929 |
| Carlos1 | Belgium | *Zea mays* | 2021 | ANVAL01 |
| Carlos2 | Belgium | *Phaseolus vulgaris* | 2021 | JANXIG01 |
| Carlos3 | Belgium | *Triticum aestivum* | 2021 | JANXIF01 |
| Carlos4 | Belgium | *Phaseolus vulgaris* | 2021 | JANVAK01 |
| Carlos5 | Belgium | *P. vulgaris* | 2021 | JANWTN01 |
| Carlos6 | Belgium | *G. max* | 2021 | JANXIE01 |
| Carlos7 | Belgium | *P. vulgaris* | 2021 | JANXID01 |
| CFBP 3417 | USA | *P. vulgaris* | 1958 | JAHEWX01 |
| CFBP 3422 | USA | *P. vulgaris* | 1956 | JAHEWY01 |
| CFBP 3423 | USA | *P. vulgaris* | 1957 | JAHEWZ01 |
| CFBP 8818 | Iran | *Solanum lycopersicum* | 2015 | JAHEWT01 |
| CFBP 8819 | Iran | *P. vulgaris* | 2014 | JAHEWS01 |
| CFBP 8820 | Iran | *Capsicum annuum* | 2015 | JAHEWR01 |
| CFBP 8821 | Iran | *Solanum melongena* | 2014 | JAHEWQ01 |
| CFBP 8822 | Iran | *C. annuum* | 2015 | JAHEWP01 |
| CFBP 8823 | Iran | *P. vulgaris* | 2015 | JAHEWO01 |
| CFBP 8824 | Iran | *S. lycopersicum* | 2015 | JAHEWN01 |
| CFBP 8825 | Iran | *S. lycopersicum* | 2015 | JAHEWM01 |
| Cff1037 | Turkey | *P. vulgaris* | 2015 | CP041259 |
| DSM 20129 | USA | *P. vulgaris* | 1990 | CP080395 |
| F30_1 | Russia | *G. max* | 2021 | CP080395 |
| LMG 3640 | Belgium | *P. vulgaris* | 2021 | JANUGT01 |
| LMG 3645 | Hungary | *P. vulgaris* | 1957 | JANVAF01 |
| LMG 3647 | Belgium | *P. vulgaris* | 2021 | JABMCF01 |
| LMG 3648 | Belgium | *Phaseolus* sp. | 2021 | JANWTK01 |

**Table S7.** Pathogenicity tests using *Curtobacterium flaccumfacien*s pv. *flaccumfaciens* isolates. +++ indicates that the isolate was able to induce symptoms on all the three replicates tested, ++ only 75% of the replicates developed symptoms, - indicates that none of the replicates tested developed symptoms.

| **Strain** | **Has plasmid pCff119?** | ***Vigna unguiculata* (cv. Red Caloona)** | ***Vigna mungo* (cv. Onyx-AU)** | ***Vigna radiata* (cv. Jade)** |
|---|---|---|---|---|
| **BRIP 70623** | Yes | +++ | +++ | +++ |
| **BRIP 70624** | No | - | - | - |
| **BRIP 70606** | Yes | +++ | +++ | +++ |
| **BRIP 70614** | Yes | +++ | ++ | +++ |
| **Cff089** | No | - | - | - |
| **DAR 58700** | No | - | - | - |
| **Mock-inoculated control** | - | - | - | - |

**Table S8.** Replicated DNA samples in whole genome sequencing of Australian *Curtobacterium flaccumfacien*s pv. *flaccumfaciens* isolates. Only the first replicate was retained for analyses and additional replicates highlighted in grey ($n$ = 11) were removed from the data set after establishing genetic distance between replicates on each isolate was zero.

| **Isolate** | **Replicate** | **Sequencing run** |
|---|---|---|
| Cff070 | 1 | 1 |
| | 2 | 1 |
| | 3 | 2 |
| Cff071 | 1 | 1 |
| | 2 | 1 |
| Cff072 | 1 | 1 |
| | 2 | 1 |
| | 3 | 2 |
| Cff192 | 1 | 1 |
| | 2 | 1 |
| Cff199 | 1 | 1 |
| | 2 | 1 |
| Cff200 | 1 | 1 |
| | 2 | 2 |
| | 3 | 2 |
| CffT14103 | 1 | 1 |
| | 2 | 2 |
| | 3 | 2 |

### References

Cingolani P et al. 2012. A program for annotating and predicting the effects of single nucleotide polymorphisms, SnpEff: SNPs in the genome of Drosophila melanogaster strain w $^{1118}$; iso-2; iso-3. Fly (Austin). 6:80–92. doi: 10.4161/fly.19695.

Jain C, Rodriguez-R LM, Phillippy AM, Konstantinidis KT, Aluru S. 2018. High throughput ANI analysis of 90K prokaryotic genomes reveals clear species boundaries. Nat. Commun. 9:5114.

Kamvar ZN, Brooks JC, GrÃ¼nwald NJ. 2015. Novel R tools for analysis of genome-wide population genetic data with emphasis on clonality. Front. Genet. 6. doi: 10.3389/fgene.2015.00208.

Kamvar ZN, Tabima JF, Grünwald NJ. 2014. *Poppr* : an R package for genetic analysis of populations with clonal, partially clonal, and/or sexual reproduction. PeerJ. 2:e281. doi: 10.7717/peerj.281.

Nei M. 1978. Estimation of average heterozygosity and genetic distance from a small number of individuals. Genetics. 89:583–590. doi: 10.1093/genetics/89.3.583.

Pritchard JK, Stephens M, Donnelly P. 2000. Inference of Population Structure Using Multilocus Genotype Data. Genetics. 155:945–959. doi: 10.1093/genetics/155.2.945.

Simpson EH. 1949. Measurement of Diversity. Nature. 163:688–688. doi: 10.1038/163688a0.

Stamatakis A. 2014. RAxML version 8: a tool for phylogenetic analysis and post-analysis of large phylogenies. Bioinformatics. 30:1312–1313. doi: 10.1093/bioinformatics/btu033.
